# A c-di-AMP-controlled glutamine synthesis pathway promotes persistence of *Staphylococcus aureus* thymidine-dependent small colony variants in the lung

**DOI:** 10.64898/2026.07.30.741254

**Authors:** Joshua P. Leeming, Shoukai Kang, Pearl P. Thakkar, Pratyusha Gorige, Devanshi R. Patel, Yuxin Pan, Oscar E. Villasana Espinosa, Daniel J. Wolter, Cara C. Boutte, Qing Tang

## Abstract

Children with cystic fibrosis (CF) commonly harbor *Staphylococcus* aureus thymidine-dependent small-colony variants (TD-SCVs), which are associated with reduced lung function and increased respiratory exacerbations. How TD-SCVs survive in the thymidine-limited CF lung is unknown. Here, we show that TD-SCVs exhibit impaired glutamine uptake and depend on c-di-AMP-regulated *de novo* glutamine synthesis for survival in the murine lung. We found that transcription of the glutamine synthetase gene *glnA* is cooperatively repressed by the transcriptional regulator GlnR, the c-di-AMP-binding protein PstA, and GlnA itself. Glutamine starvation elevates c-di-AMP levels, relieving repression by this ternary complex and promoting glutamine synthesis. Reducing c-di-AMP levels causes a profound growth defect in TD-SCVs under low-thymidine conditions, which is rescued by *glnA* overexpression. Moreover, pharmacological inhibition of GlnA markedly impairs TD-SCV growth in murine lung. These findings elucidate the molecular mechanism underlying *S. aureus* TD-SCV survival during infection and identify glutamine synthesis as a promising therapeutic target for treating infections caused by antifolate-resistant bacteria.

## Main

*Staphylococcus aureus* is the leading bacterial cause of death in 135 countries and is associated with over one million deaths worldwide a year^1^. Individuals with cystic fibrosis (CF), who have thickened mucus and dysregulated lung immune microenvironments due to mutations in the cystic fibrosis transmembrane conductance regulator (*CFTR)* gene, are highly susceptible to *S. aureus* lung infections^2^. Over the past decades, *S. aureus* has been the most prevalent organism infecting the respiratory tract of children with CF and remains the second most prevalent organism in adults with CF, after *Pseudomonas aeruginosa*^3^. In children with CF under two years of age, more than 50% are colonized with *S. aureus*, and prevalence peaks at nearly 80% during early adolescence^4^. With advances in antimicrobial therapy, the prevalence of *P. aeruginosa* in CF lungs has continued to decline, whereas *S. aureus* has remained stable and is the most prevalent respiratory pathogen in CF in recent years^5^.

Moreover, individuals with CF commonly have complex polymicrobial infections coexisting with *S. aureus*. Anti-folates, such as the sulfamethoxazole-trimethoprim antibiotic combination, are commonly used to treat infections caused by *Burkholderia cepacia* complex, *Stenotrophomonas maltophilia*, and methicillin-resistant *S. aureus* in CF patients^6–8^. However, extensive anti-folate treatment leads to the emergence of *S. aureus* thymidine-dependent small colony variants (TD-SCVs) in respiratory tract^9,10^, particularly in CF children^11,12^. Multiple studies of CF patients worldwide have reported that *S. aureus* TD-SCV colonization is associated with reduced lung function and an increased risk of respiratory exacerbations compared to either normal colony (NC) *S. aureus* or SCVs of other small colony genotypes^9,11,12^. These mutants are thymidine auxotrophs resulting from inactivating mutations in thymidylate synthase (ThyA) and rely on exogenous thymidine for growth^13^. Thymidine concentrations in the sputum of CF patients vary widely, ranging from 0.1 to 38 μg/ml^14^, whereas *S. aureus* TD-SCVs require a minimum of 2.5 μg/ml thymidine for normal growth^15^. These observations suggest that many patients may not have sufficient thymidine levels to fully support TD-SCV growth. In addition, despite the overall reported range, local microenvironments within the thick CF mucus may remain thymidine limited. It remains unclear how *S. aureus* TD-SCVs survive in CF lungs with low thymidine levels and cause long-term infections. This knowledge gap has significantly hindered the development of therapeutic strategies for controlling chronic and recurrent infections caused by TD-SCVs.

Our previous studies revealed that *S. aureus* TD-SCVs produce elevated levels of cyclic di-AMP (c-di-AMP) during infection, which activates the immune mediator STING, leading to increased airway inflammation that potentially contributes to severe lung outcomes in CF patients^15^. Despite this progress, it has remained unclear whether elevated c-di-AMP production is detrimental or beneficial for the survival of *S. aureus* TD-SCVs during infection. In addition to serving as an inducer of eukaryotic immune responses^16^, c-di-AMP functions as a second messenger in more than 11,000 bacterial and archaeal species^17^, regulating essential cellular processes such as carbon metabolism, osmotic homeostasis, cell wall synthesis, and DNA integrity through allosteric regulation of various RNA riboswitches and proteins^18,19^. In *S. aureus*, c-di-AMP production changes in response to both thymidine and glutamine concentrations^15,20^. Bacteria synthesize dUMP from glutamine through a series of pyrimidine synthesis enzymes, and dUMP is subsequently converted to dTMP by ThyA for DNA synthesis. In *S. aureus* TD-SCVs, in which ThyA is inactivated, cells rely on exogenous thymidine for dTMP production. These results suggest a potential role for c-di-AMP in coordinating glutamine and thymidine availability to regulate TD-SCV survival.

Glutamine is a metabolite of central importance to nitrogen metabolism and bacterial physiology. It is required for protein synthesis, acts as a nitrogen donor during the biosynthesis of various nitrogen-containing compounds, and serves as a signaling molecule^21^. *Bacillus subtilis* has been used as a primary model for studying glutamine metabolism in Gram-positive bacteria. In *B. subtilis*, the transcription of the glutamine synthetase gene *glnA* is activated by TnrA and repressed by GlnR in response to glutamine levels. In the absence of glutamine, TnrA induces *glnA* expression, resulting in glutamine biosynthesis by GlnA from glutamate and ammonium^22^. In the presence of glutamine, GlnA acts as a chaperone that deactivates TnrA while also binding to GlnR and strongly promoting repression of *glnA* expression^23^. Furthermore, the PII family protein GlnK promotes *glnA* expression by stabilizing TnrA under low-glutamine conditions, while inhibiting GlnA synthesis in the presence of glutamine by blocking ammonium transport^22,24^. *S. aureus*, like most Firmicutes, has GlnA and GlnR but not TnrA^25,26^, leaving it unclear how these bacteria sense glutamine concentrations and regulate glutamine synthesis. In addition, *S. aureus* has only a truncated GlnK; instead, it encodes an additional PII family protein, PstA. PstA is structurally distinct from GlnK and binds c-di-AMP^27^, although its downstream binding targets remain unidentified. These observations further indicate that c-di-AMP-PstA signaling may compensate for the loss of TnrA- and GlnK-mediated regulation of glutamine metabolism in *S. aureus*.

In this study, we revealed a previously unappreciated *S. aureus* glutamine synthesis regulatory circuitry mediated by c-di-AMP. In the presence of glutamine, c-di-AMP production is inhibited, and GlnA and PstA cooperatively enhance the DNA-binding ability of GlnR, resulting in suppression of glutamine synthesis genes. In the absence of glutamine, elevated c-di-AMP abolishes this cooperative enhancement by PstA and GlnA, leading to increased *glnA* expression and glutamine synthesis. Furthermore, we found that *de novo* glutamine synthesis, rather than exogenous glutamine uptake, is required for *S. aureus* TD-SCV growth under low-thymidine conditions and in murine lung. Our previous study found that *S. aureus* TD-SCVs produce elevated levels of c-di-AMP under low-thymidine conditions^15^. Here, we show that reducing c-di-AMP levels in *S. aureus* TD-SCVs leads to a dramatic growth defect under low-thymidine conditions, and that this defect can be fully rescued by *glnA* overexpression. More importantly, pharmacological inhibition of GlnA results in growth defects of *S. aureus* TD-SCVs in both macrophages and mice. Our study therefore reveals how glutamine metabolism promotes the survival and pathogenesis of *S. aureus* and provides a mechanistic explanation for how *S. aureus* TD-SCVs survive thymidine starvation during infection. Our findings also highlight glutamine synthesis as a potential therapeutic target for combating chronic and recurrent TD-SCV infections.

## Results

### Transcriptional repressor GlnR regulates *S. aureus* glutamine synthesis

In *S. aureus*, glutamine is synthesized by GlnA from glutamate and ammonium (NH4^+^). Ammonium can be imported by the ammonium transporter AmtB (**Fig. 1a**). Given the critical role of glutamine in central metabolism, its metabolism must be precisely regulated. However, the regulation of glutamine metabolism in *S. aureus* remains unclear.

**Figure 1.**
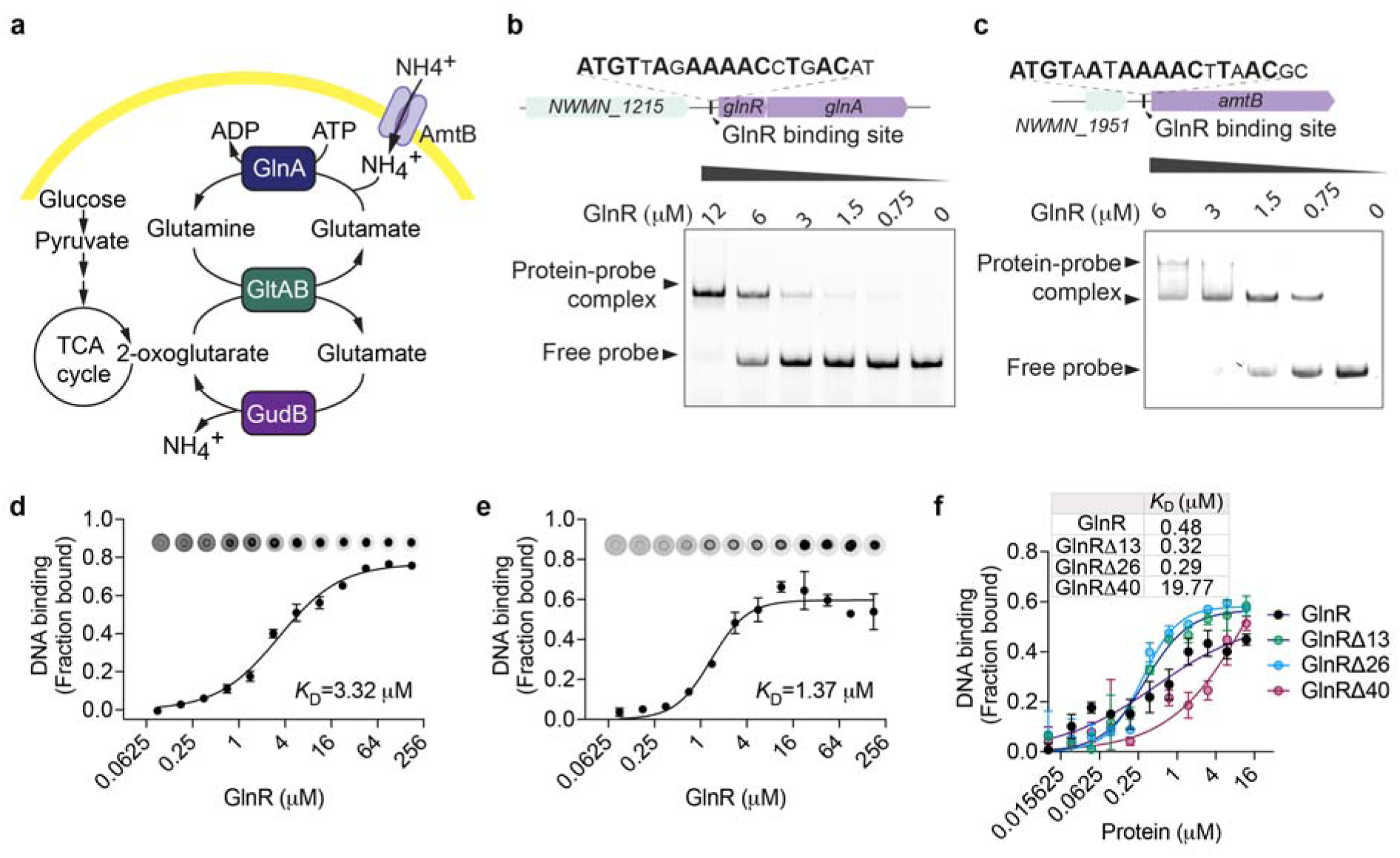
GlnR binds to the promoter region of glutamine synthesis genes. **a**, Schematic diagram of *de novo* glutamine synthesis. **b**-**c**, EMSA revealed the *in vitro* binding of GlnR to the 59-nt promoter region of the *glnRA* operon (**b**) or the 60-nt promoter region of the *amtB* gene (**c**). **d-e**, DRaCALA analysis of GlnR DNA-binding ability to the *glnRA* (**d**) or *amtB* (**e**) promoter. The upper panel shows DRaCALA images, while the lower panel presents the DNA fraction bound to GlnR in duplicates. **f**, DRaCALA analysis of the DNA-binding abilities of GlnR and its truncated mutants to the *glnRA* promoter. 0.2 μM 5’-FAM-DNA probe was incubated with GlnR in both EMSA and DRaCALA assays. The fraction bound of DRaCALA was calculated by normalizing the intensity of the protein-DNA complex to the background intensity. For panels **d-f**, mean values of technical duplicates are plotted, and error bars indicate ± SD, and The *K*_D_ was calculated using GraphPad Prism with the “specific binding with Hill slope” model.

Our bioinformatic studies identified conserved binding motifs of the transcriptional repressor GlnR in the promoter regions of the *glnA-glnR* operon, as well as the *amtB* gene in *S. aureus*. Furthermore, we demonstrated that GlnR binds to these promoters, as assessed by <u>E</u>lectrophoretic <u>M</u>obility <u>S</u>hift <u>A</u>ssay (EMSA) assays. Binding to a high concentration of GlnR led to slower migration of the DNA probes on the EMSA gel (**Fig. 1b, c**). Similar results were obtained using <u>D</u>ifferential <u>Ra</u>dial <u>C</u>apillary <u>A</u>ction of <u>L</u>igand assay (DRaCALA) by incubating 5’-FAM-labeled DNA probes with GlnR and observing the formation of DNA-probe complexes on the nitrocellulose membrane (**Fig. 1d, e**).

GlnRs in Firmicutes contain a conserved N-terminal DNA-binding domain and a flexible C-terminal domain (**Extended Data Fig. 1a**). The C-terminal domain of *B. subtilis* GlnR autoinhibits its dimerization, which is required for DNA binding, and the apo-GlnR exists primarily as a monomer. In contrast, *S. aureus* GlnR predominantly forms a dimer in solution, regardless of DNA supplementation (**Extended Data Fig. 2a, b**). To investigate the function of the C-terminal domain of *S. aureus* GlnR, His_6_-tagged GlnR and its C-terminal truncation mutants, GlnRΔ13, GlnRΔ26, and GlnRΔ40, were purified (**Extended Data Fig. 1b**), and their DNA-binding abilities were analyzed using DRaCALA. We found that both GlnRΔ13 and GlnRΔ26 exhibited a moderate increase in DNA-binding ability compared with wild-type GlnR, whereas GlnRΔ40, which lacks the entire C-terminal regulatory domain, displayed reduced DNA-binding activity (**Fig. 1f**). These data indicate that although *S. aureus* GlnR possesses an autoinhibitory mechanism mediated by its 26-amino acid C-terminus, this inhibition is not robust, suggesting the presence of additional regulatory mechanisms controlling GlnR activity.

### Glutamine synthetase GlnA strongly enhances the DNA-binding of GlnR

Previous studies in *B. subtilis* have shown that GlnA is a bifunctional protein that functions as an enzyme for glutamine synthesis under low cellular glutamine concentrations and acts as a chaperone for GlnR, enhancing its DNA-binding ability to repress glutamine synthesis gene expression under high glutamine concentrations^23^. Consistently, we found that *S. aureus* GlnA strongly enhances the DNA-binding capacity of GlnR (**Fig. 2a**). Similar to the GlnA-GlnR interaction in *B. subtilis*^23^, we did not observe a supershifted band in the EMSA assay, indicating that *S. aureus* GlnA may regulate GlnR function as a chaperone-like factor that does not need to form a stable complex with GlnR. GlnR proteins in Firmicutes contain a conserved N-terminal DNA-binding domain and a C-terminal regulatory domain with variable amino acid sequences^28^ (**Extended Data Fig. 1a**). To investigate the role of the C-terminal regulatory domain of GlnR in mediating its interaction with GlnA, we examined the effect of GlnA on the DNA-binding ability of full-length GlnR and its C-terminal truncated mutants. Deletion of 40 amino acids from the C-terminal domain of GlnR (GlnRΔ40) largely retained its DNA-binding ability (**Fig. 1f**); however, it abolished the interaction with GlnA (**Fig. 2b** and **Extended Data Fig. 3**).

**Figure 2.**
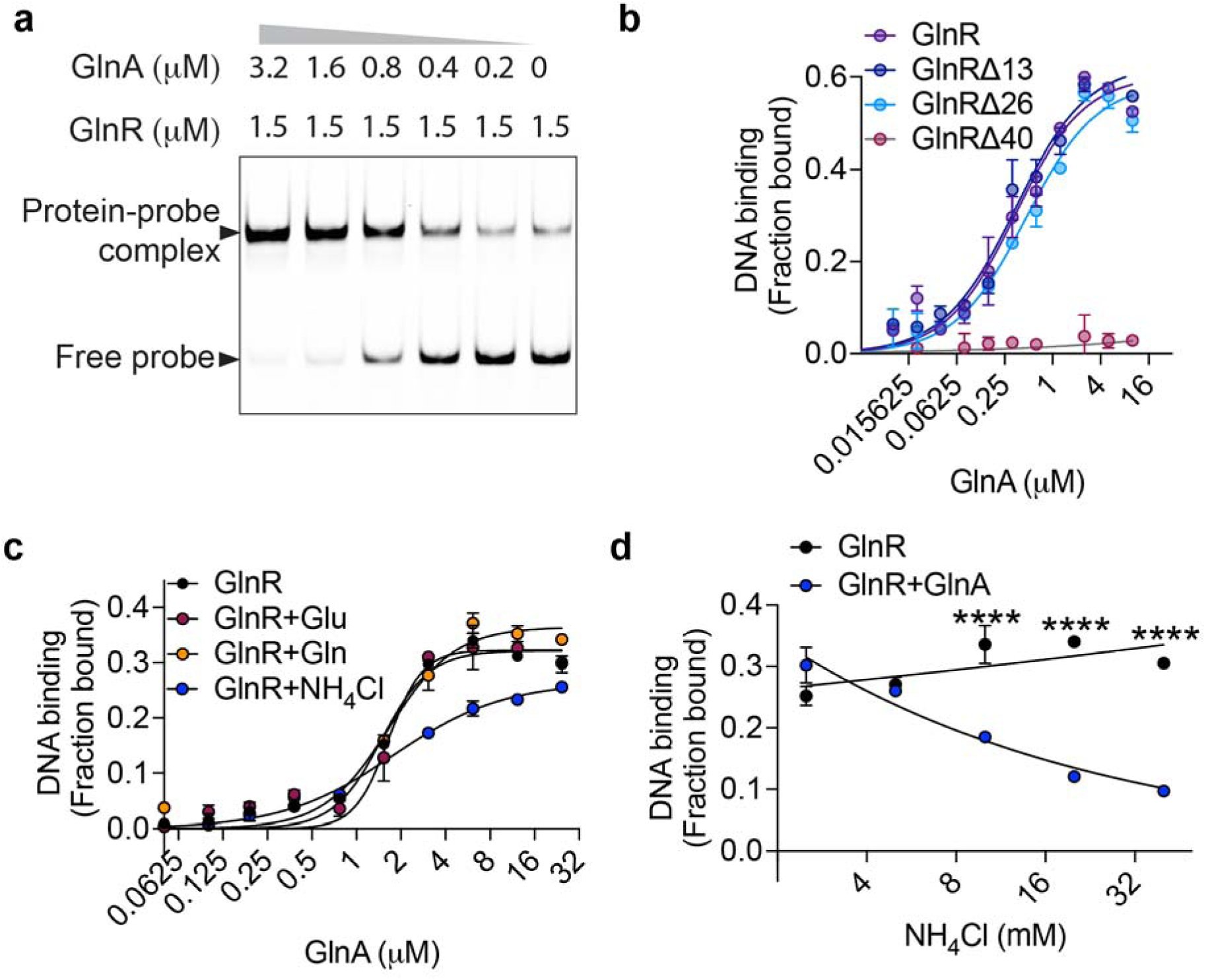
GlnA promotes the DNA-binding ability of GlnR independent of glutamine. **a**, EMSA analysis of GlnR DNA-binding in the presence of increasing concentrations of GlnA. **b**, DRaCALA analysis of GlnR and its truncated mutants binding to DNA in the presence of increasing concentrations of GlnA. 0.25 μM GlnR, 0.15 μM GlnRΔ13, 0.125 μM GlnRΔ26 and 4 μM GlnRΔ26 was used. **c**, DRaCALA analysis of the impact of GlnA on GlnR DNA-binding in the presence of 20 mM glutamine, glutamate, or NH_4_Cl. 0.5 μM GlnR was used. **d**, DRaCALA analysis of the impact of GlnA on GlnR DNA-binding in the presence of increasing concentrations of NH_4_Cl. 0.5 μM GlnR and 2 μM glnA was used. 0.2 μM 5’-FAM-DNA probe (*glnRA* promoter region) was used in both EMSA and DRaCALA assays. GlnA protein used in panel **c** and **d** was purified using urea denaturation and renaturation to deplete its potential bound ligands. For panels **d-f**, mean values of technical duplicates are plotted, and error bars indicate ± SD. *P* values were calculated using two-way ANOVA. Asterisks indicate that differences are statistically significant (****, *P* < 0.0001), and “ns” indicates no significant difference.

In contrast to *B. subtilis*, the promotion of GlnR DNA-binding by GlnA was inhibited by ammonium, whereas supplementation with glutamine or glutamate had no effect (**Fig. 2c**). Ammonium appears to affect GlnR DNA binding indirectly through GlnA rather than acting directly on GlnR, and a minimum concentration of 10 mM ammonium was required for significant inhibition (**Fig. 2d**). These data indicate that GlnA interacts with the C-terminal domain of GlnR and promotes its DNA-binding, which is feedback-inhibited by high concentrations of ammonium. In humans, physiological serum ammonia levels are typically very low (10-50 µM)^29^, suggesting that *S. aureus* may possess additional feedback mechanisms that are more sensitive for detecting nitrogen sources.

### The PII family protein PstA promotes the DNA-binding of GlnR in *S. aureus*

PII family signal transduction proteins are among the most widely distributed signaling proteins in bacteria, with a primary role in regulating nitrogen metabolism^30–32^. GlnB and GlnK are members of the PII signal transduction family. GlnB is found in Proteobacteria and Cyanobacteria, whereas GlnK is conserved in Proteobacteria and Firmicutes^30^. In response to nitrogen availability, GlnB and GlnK undergo posttranslational modification and directly interact with the ammonium transporter (AmtB) and glutamine synthetase (GlnA), thereby regulating the glutamine synthesis^21,30^. *S. aureus* lacks both GlnB and GlnK but contains a distinct PII protein, PstA. PstA is a conserved c-di-AMP-binding protein found in several pathogenic Firmicutes, including *S. aureus*, *Listeria monocytogenes*, *Clostridium botulinum*, and *Enterococcus faecalis*^33–35^.

PII signaling proteins form regulatory protein-protein interactions that are inhibited upon metabolite binding. Using double-reference-subtracted biolayer interferometry (BLI), we detected a concentration-dependent interaction between PstA and GlnR, with an apparent *K*_D_ of approximately 60 μM. This interaction was enhanced approximately sevenfold in the presence of c-di-AMP, with the apparent *K*_D_ decreasing to approximately 8 μM (**Fig. 3a**). We next asked whether PstA influences GlnR DNA binding. In EMSA assays, the PstA-GlnR interaction enhanced the DNA-binding ability of GlnR (**Fig. 3b**), although this effect was less pronounced than that of GlnA (**Fig. 2a**). We did not observe a supershifted complex in EMSA, suggesting that PstA may enhance GlnR DNA binding through a weak or transient interaction that is not maintained during electrophoresis. We performed BLI assays using biotinylated *glnAR* promoter DNA captured on a streptavidin biosensor and consistently observed that PstA enhances the DNA-binding activity of GlnR (**Fig. 3c**). Consistent with the GlnA-GlnR interaction, the C-terminal regulatory region (residues 26-40) is required for PstA-dependent enhancement of DNA binding (**Fig. 3d, e**). Although c-di-AMP promotes the PstA-GlnR interaction detected by BLI (**Fig. 3a**), it caused only a modest, non-significant enhancement of GlnR DNA binding in the presence of PstA (**Fig. 3f**). These data suggest that PstA enhances GlnR DNA binding, whereas the major regulatory effect of c-di-AMP may require additional components of the GlnR regulatory complex.

**Figure 3.**
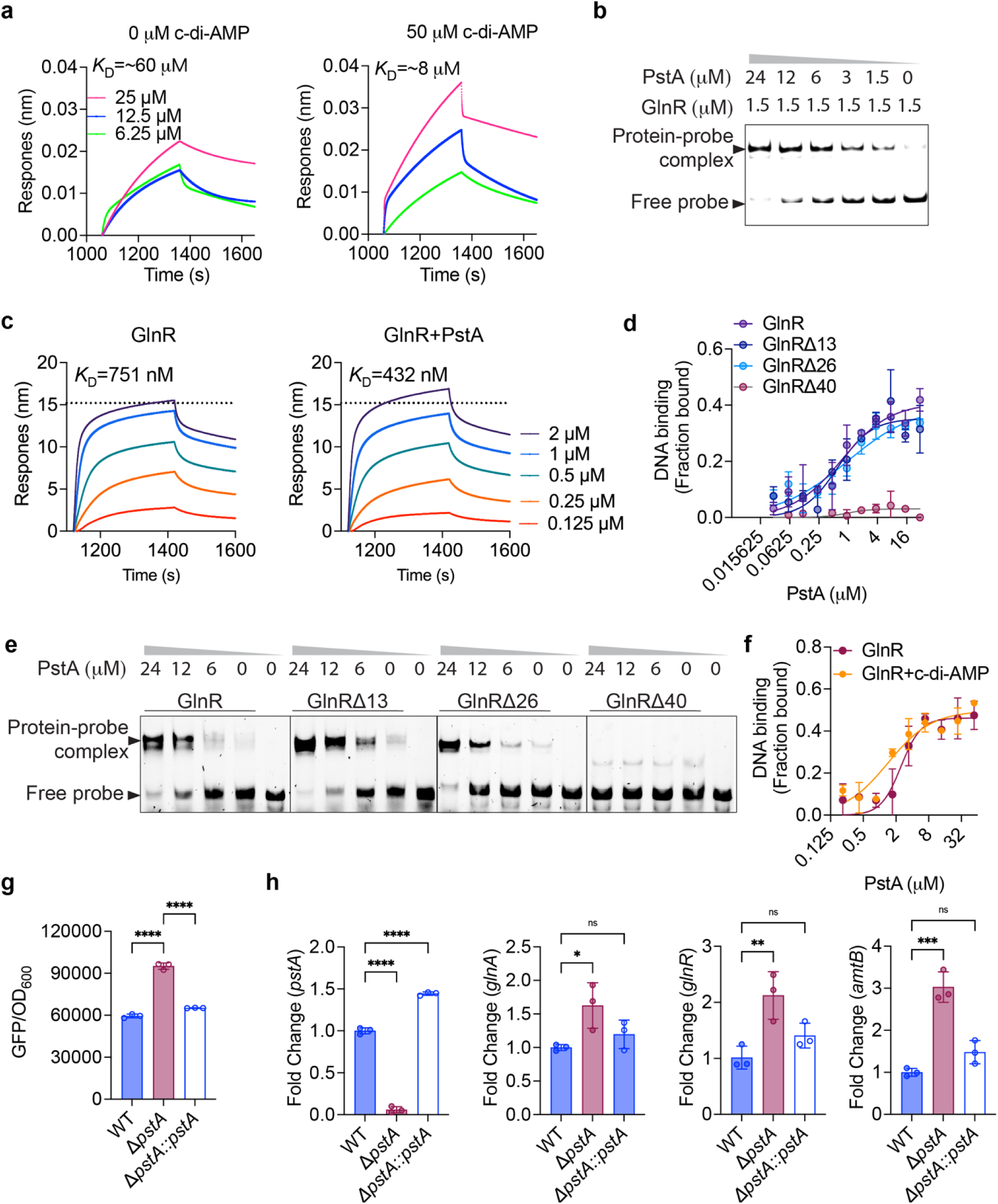
PstA promotes the DNA binding ability of GlnR. **a**, *In vitro* binding of GlnR to PstA analyzed by BLI. Biotinylated-PstA was pre-conjugated with the streptavidin sensor, and an increasing concentration of the GlnR was tested for protein-protein interactions, with or without c-di-AMP. **b**, EMSA analysis of GlnR DNA-binding in the presence of increasing concentrations of the PstA. **c**, *In vitro* binding of GlnR to *glnRA* promoter examined in the presence of PstA by BLI. The 5′-biotinylated glnRA promoter fragment (59-nt) was pre-conjugated to the streptavidin sensor, and increasing concentrations of GlnR, in the presence or absence of 50 μM PstA, were tested for DNA binding. **d-e**, DRaCALA (**d**) or EMSA (**e**) analysis of GlnR and its truncated mutants binding to DNA in the presence of increasing concentrations of PstA. 0.25 μM GlnR, 0.15 μM GlnRΔ13, 0.125 μM GlnRΔ26 and 4 μM GlnRΔ26 was used. **f**, DRaCALA analysis of the impact of PstA on GlnR DNA-binding in the presence or absence of 25 μM c-di-AMP. **g**, The *glnAR* promoter activity in the *S. aureus* Newman strains grown in BHI was assessed by measuring GFP intensity driven by the *glnAR* promoter, with the data normalized to OD_600_. **h**, The *pstA*, *glnR*, *glnA*, and *amtB* gene expression in the *S. aureus* Newman strains grown in BHI as measured by qRT-PCR. The mRNA levels were normalized to those of the WT strain using 16S rRNA as the internal control. For panels **d** and **f**, mean values of technical duplicates are plotted. For panels **g** and **h**, mean values of technical replicates are plotted, and error bars indicate ± SD. *P* values were calculated using two-way ANOVA. Asterisks indicate that differences are statistically significant (*, *P* < 0.05; **, *P* < 0.01; ***, *P* < 0.001, ****, *P* < 0.0001), and “ns” indicates no significant difference.

To determine whether PstA regulates GlnR activity in cells, we generated a *pstA* knockout (Δ*pstA*) strain and a complementation strain (Δ*pstA*::*pstA*), and measured *glnAR* promoter activity by assessing GFP expression driven by the *glnAR* promoter in these strains. We observed significantly higher *glnAR* promoter activity in the Δ*pstA* strain compared to both the WT and Δ*pstA*::*pstA* strains when grown in BHI (**Fig. 3g**). Consistently, qRT-PCR showed increased expression of GlnR-regulated genes in Δ*pstA*, including *glnR*, *glnA* and *amtB* (**Fig. 3h**). Together, these results support a model in which PstA functions as a co-regulator that promotes GlnR-dependent DNA binding and transcriptional repression of glutamine metabolism genes in *S. aureus*.

### C-di-AMP abolishes the cooperative enhancement of GlnR DNA-binding mediated by PstA and GlnA

Our data has indicated that GlnA and PstA independently promote the DNA-binding ability of GlnR; however, it is unclear how these proteins coordinate with each other in *S. aureus*. We found that a low concentration of PstA could promote the DNA-binding ability of GlnR in the presence of GlnA (**Fig. 4a**), suggesting an additive role for PstA and GlnA in enhancing GlnR DNA binding. C-di-AMP bound only to PstA, but not to GlnR or GlnA, when we tested binding of these proteins to ^32^P-c-di-AMP using DRaCALA (**Fig. 4b**). Notably, c-di-AMP abolished the additive enhancement of GlnR’s DNA-binding by PstA and GlnA, while having no effect on GlnR in the presence of either GlnA or PstA alone (**Fig. 4c**). To further investigate the specificity of c-di-AMP in regulating the interactions among PstA, GlnR, and GlnA, we generated a c-di-AMP-blind mutant of PstA by mutating the conserved c-di-AMP-binding motif GGFL to alanines. The c-di-AMP binding ability of PstA_GGFL-AAAA_ is abolished as assessed by DRaCALA (**Fig. 4d**), yet it still promotes the DNA-binding activity of GlnR (**Fig. 4e**). Notably, the DNA-binding ability of GlnR remained affected by increasing c-di-AMP levels in the presence of GlnA and PstA_GGFL-AAAA_ (**Fig. 4f**). These data suggest that c-di-AMP, via PstA, could promote expression of glutamine metabolic genes through modulating the DNA-binding state of the GlnA-PstA-GlnR complex.

**Figure 4.**
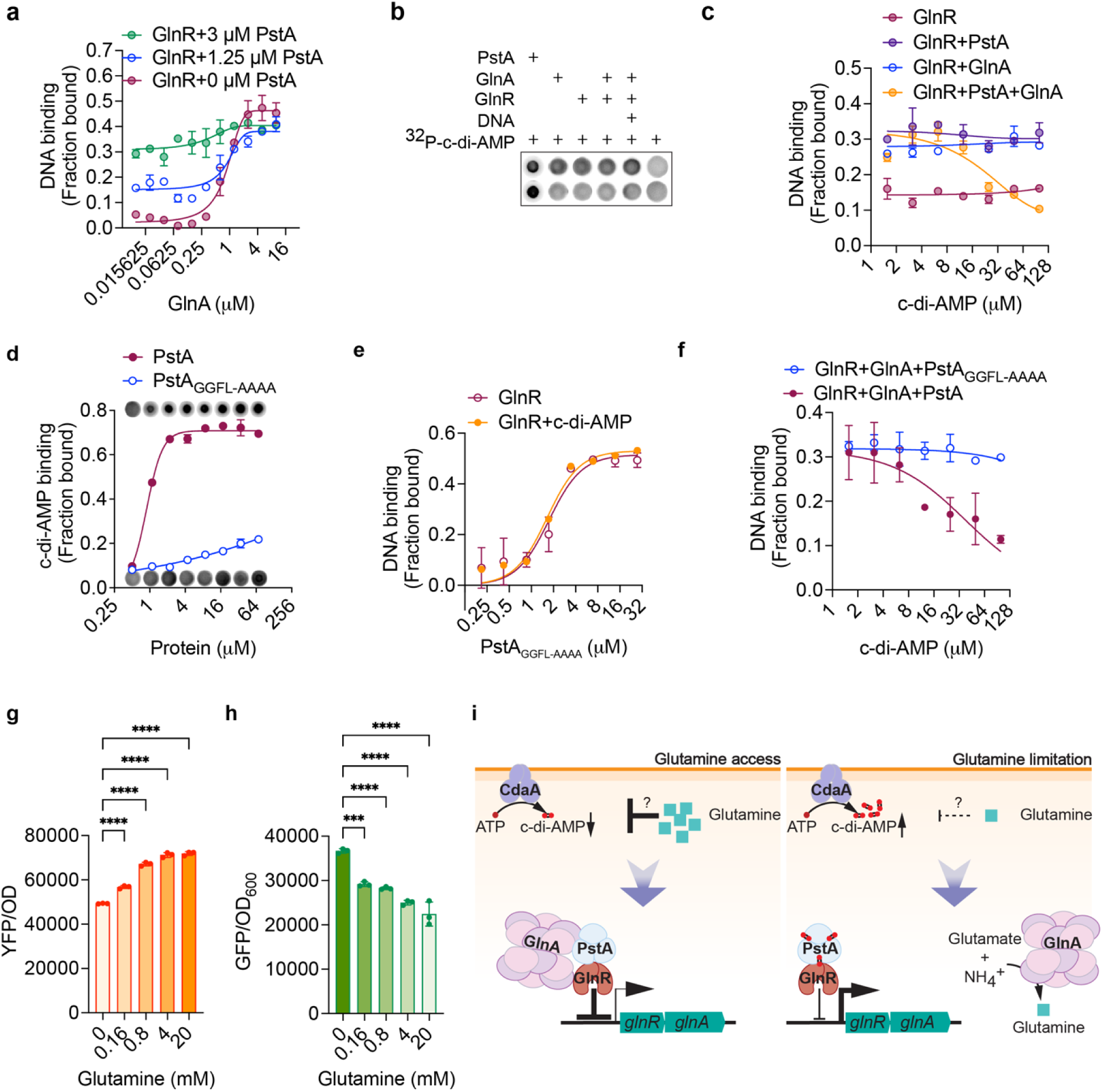
C-di-AMP promotes the dissociation of GlnA from GlnR by PstA. **a**, DRaCALA analysis of GlnR DNA-binding in the presence of both PstA and GlnA. 0.5 μM GlnR was used. **b**, *In vitro* binding of ^32^P-c-di-AMP to PstA, GlnR, or GlnA analyzed by DRaCALA. **c**, DRaCALA analysis of GlnR DNA-binding in the presence of PstA, GlnA, or both, with increasing concentrations of c-di-AMP. 0.5 μM GlnR, 0.625 μM GlnA and 3 μM PstA was used. **d**, *In vitro* binding of ^32^P-c-di-AMP to PstA and PstA_GGFL-AAAA_ analyzed by DRaCALA. **e**, DRaCALA analysis of the impact of PstA_GGFL-AAAA_ on GlnR DNA-binding in the presence or absence of 25 μM c-di-AMP. **f**, DRaCALA analysis of the impact of PstA or PstA_GGFL-AAAA_ on GlnR DNA binding in the presence of increasing concentrations of c-di-AMP. **g**, Intracellular c-di-AMP concentration quantified by c-di-AMP-reporter assay. **h**, The *glnAR* promoter activity in the *S. aureus* Newman strains grown in SCFM with increasing concentration of glutamine supplementation was assessed by measuring GFP intensity driven by the *glnAR* promoter, with the data normalized to OD_600._ **i**, Proposed schematic diagram of glutamine synthesis regulation in *S. aureus*. High glutamine inhibits c-di-AMP production, allowing PstA and GlnA to enhance GlnR DNA binding and repress glutamine synthesis genes. Low glutamine elevates c-di-AMP, disrupts this interaction, derepresses these genes, and frees GlnA to catalyze glutamine synthesis. For panels **a**, **c**, **d**, **e**, and **f**, mean values of technical duplicates are plotted; for panels **g** and **h**, biological triplicates are plotted. For all panels, error bars indicate ± SD. *P* values were calculated using two-way ANOVA. Asterisks indicate that differences are statistically significant (*, *P* < 0.05; **, *P* < 0.01; ***, *P* < 0.001, ****, *P* < 0.0001), and “ns” indicates no significant difference.

To investigate whether *S. aureus* alters c-di-AMP production in response to glutamine concentrations, thereby providing feedback regulation of glutamine synthesis, we quantified the intracellular c-di-AMP concentration in *S. aureus* using a c-di-AMP biosensor in which YFP expression is driven by a c-di-AMP riboswitch, and c-di-AMP binding leads to repression of YFP expression^36^. Consistent with previous studies^7^, we found that intracellular c-di-AMP production in *S. aureus* is inhibited by glutamine supplementation (**Fig. 4g**). Given that c-di-AMP relieves the transcriptional repression by GlnR (**Fig. 4c**), we hypothesize that decreasing c-di-AMP will lead to enhanced DNA binding and decreased promoter activity. Consistent with our hypothesis, we observed that glutamine supplementation, which decreases c-di-AMP concentration, dampens *glnAR* promoter activity in a concentration-dependent manner when *S. aureus* is grown in synthetic CF sputum medium (SCFM) (**Fig. 4h**).

Taken together, these observations support a model in which high glutamine concentrations inhibit intracellular c-di-AMP production, allowing PstA and GlnA to additively repress transcription of glutamine synthesis genes through GlnR (**Fig. 4i**, left panel). In contrast, when glutamine concentrations are low, c-di-AMP levels increase, disrupting the interaction among PstA, GlnA, and GlnR, thereby relieving the transcriptional repression of glutamine synthesis genes. The released GlnA then functions enzymatically in glutamine synthesis (**Fig. 4i**, right panel). Thus, this model enables both transcriptional and post-translational activation of glutamine metabolism through c-di-AMP.

### *De novo* glutamine synthesis promotes the growth of *S. aureus* TD-SCV in low thymidine conditions

*S. aureus* TD-SCVs produce elevated levels of c-di-AMP under low-thymidine conditions^15^. We have demonstrated that increased c-di-AMP promotes glutamine metabolism through PstA (**Fig. 4c, f**). To investigate how elevated c-di-AMP affects the survival of *S. aureus* TD-SCVs and whether the effects are attributable to enhanced glutamine synthesis, we generated a laboratory *S. aureus* TD-SCV strain by knocking out the *thyA* gene in *S. aureus* Newman strain. Though a whole cell proteomics analysis, we observed significant downregulation of enzymes in the purine synthesis pathway, while both the pyrimidine synthesis and DNA repair pathways were significantly upregulated in the Δ*thyA* strain grown in low thymidine (1.25 μg/ml) (**Fig. 5a, b**). Under this condition, Δ*thyA* lacks sufficient thymidine for growth, leading to upregulation of *de novo* pyrimidine synthesis and accumulation of dUTP (**Fig. 5c**). dUTP can be incorporated into DNA to support replication and bacterial growth, although this may increase the mutation rate and lead to multidrug resistance^37^, as observed in *S. aureus* TD-SCVs^38^. *S. aureus* uses glutamine for pyrimidine synthesis (**Fig. 5c**); however, the glutamine transporters GlnP and GlnQ are significantly downregulated in Δ*thyA* when grown under low-thymidine conditions (**Fig. 5d**), and the glutamine transporter AlsT was not detected by whole cell proteomics in either the WT or Δ*thyA* strains. To measure glutamine uptake, we grew WT and Δ*thyA* in BHI supplemented with ³H-glutamine, with or without 20 mM unlabeled glutamine. We found that the intracellular ³H-glutamine level in the Δ*thyA* strain was comparable to that of the WT strain when grown with high concentration of thymidine (5 μg/mL), whereas it was significantly lower than that of the WT strain under low-thymidine conditions (1.25 μg/mL). As expected, supplementation with unlabeled glutamine competitively inhibited ³H-glutamine uptake, resulting in similarly low intracellular ³H-glutamine levels in both strains irrespective of the thymidine concentration (**Fig. 5e**). These data indicate that glutamine uptake by Δ*thyA* is impaired under low-thymidine conditions.

**Figure 5.**
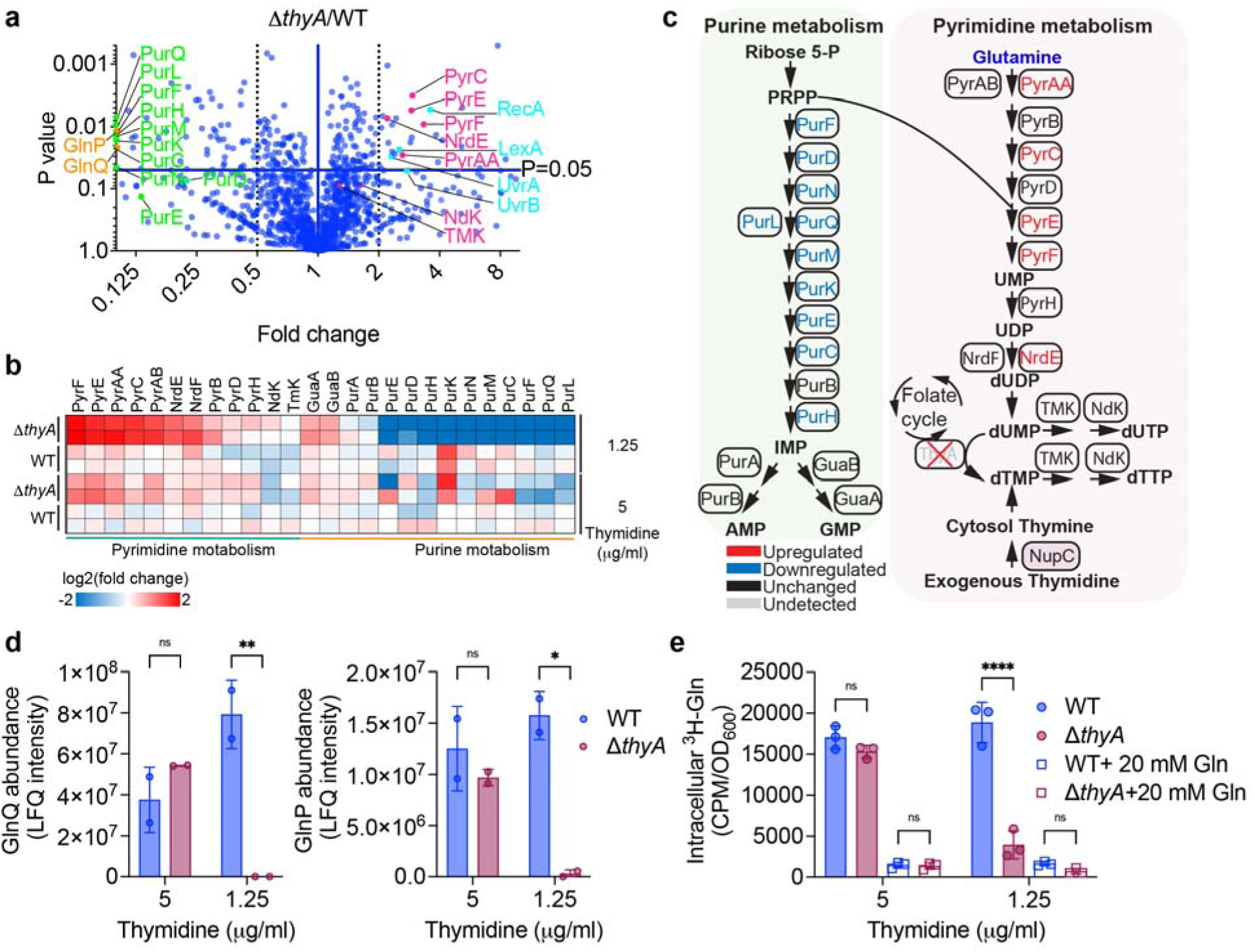
Thymidine starvation promotes pyrimidine synthesis in *S. aureus* TD-SCVs while inhibiting glutamine uptake. **a**, Whole-cell proteomic analysis of WT and Δ*thyA S. aureus* strains grown in 1.25 μg/ml thymidine. Volcano plots depict the fold change of each protein, and the *P*-value was calculated using an unpaired t-test. Magenta points represent pyrimidine synthesis proteins; green points represent purine synthesis proteins; orange points represent glutamine transport proteins; ice blue points represent DNA repair proteins. **b**, Heat map of relative protein abundance in WT and Δ*thyA* strains grown in 5 or 1.25 μg/mL thymidine, determined by whole-cell proteomic analysis. Log_2_ ratios of protein abundance for each growth condition are shown relative to WT grown in 5 μg/mL thymidine. **c**, Schematic diagram of *de novo* purine and pyrimidine synthesis. Proteins upregulated or downregulated in Δ*thyA* relative to the WT strain grown in 1.25 μg/mL thymidine are indicated in red and blue, respectively. Proteins that remain unchanged are indicated in black, and those with undetected expression levels are indicated in brown. **d**, Protein abundance of glutamine uptake proteins determined by whole-cell proteomic analysis. **e**, ³H-glutamine uptake by *S. aureus* WT and Δ*thyA* strains grown in different thymidine concentrations. *P* values were calculated using two-way ANOVA analysis; Asterisks indicate that differences are statistically significant (*, *P* < 0.05; **, *P* < 0.01, ****, *P* < 0.0001), and “ns” indicates no significant difference.

To investigate whether *de novo* glutamine synthesis promotes the growth of Δ*thyA*, we overexpressed *glnA* in both WT and Δ*thyA* strains (**Extended Data Fig. 4**). Notably, both glutamine supplementation and GlnA overexpression promoted the growth of WT in SCFM regardless of thymidine concentration (**Fig. 6a**) and enhanced the growth of Δ*thyA* at high thymidine levels (5 μg/mL) (**Fig. 6b**, left panel). However, overexpression of GlnA in Δ*thyA* restored its growth defect under low-thymidine conditions (1.25 μg/mL), whereas glutamine supplementation had no effect on the growth of Δ*thyA* (**Fig. 6b**, right panel). We observed the same phenotype upon overexpression of *glnA* in two *S. aureus* TD-SCV clinical isolates from CF children (**Fig. 6c, d**). These data support our hypothesis that the glutamine used for pyrimidine synthesis in Δ*thyA* grown in low thymidine conditions is derived from *de novo* synthesis rather than uptake. Furthermore, we found that decreasing c-di-AMP levels in Δ*thyA* by overexpressing the phosphodiesterase PdeA led to a growth defect in Δ*thyA* under low-thymidine conditions (**Fig. 6e**). This growth defect could be rescued by GlnA overexpression, further indicating that c-di-AMP promotes the growth of TD-SCVs under low-thymidine conditions by enhancing glutamine synthesis.

**Figure 6.**
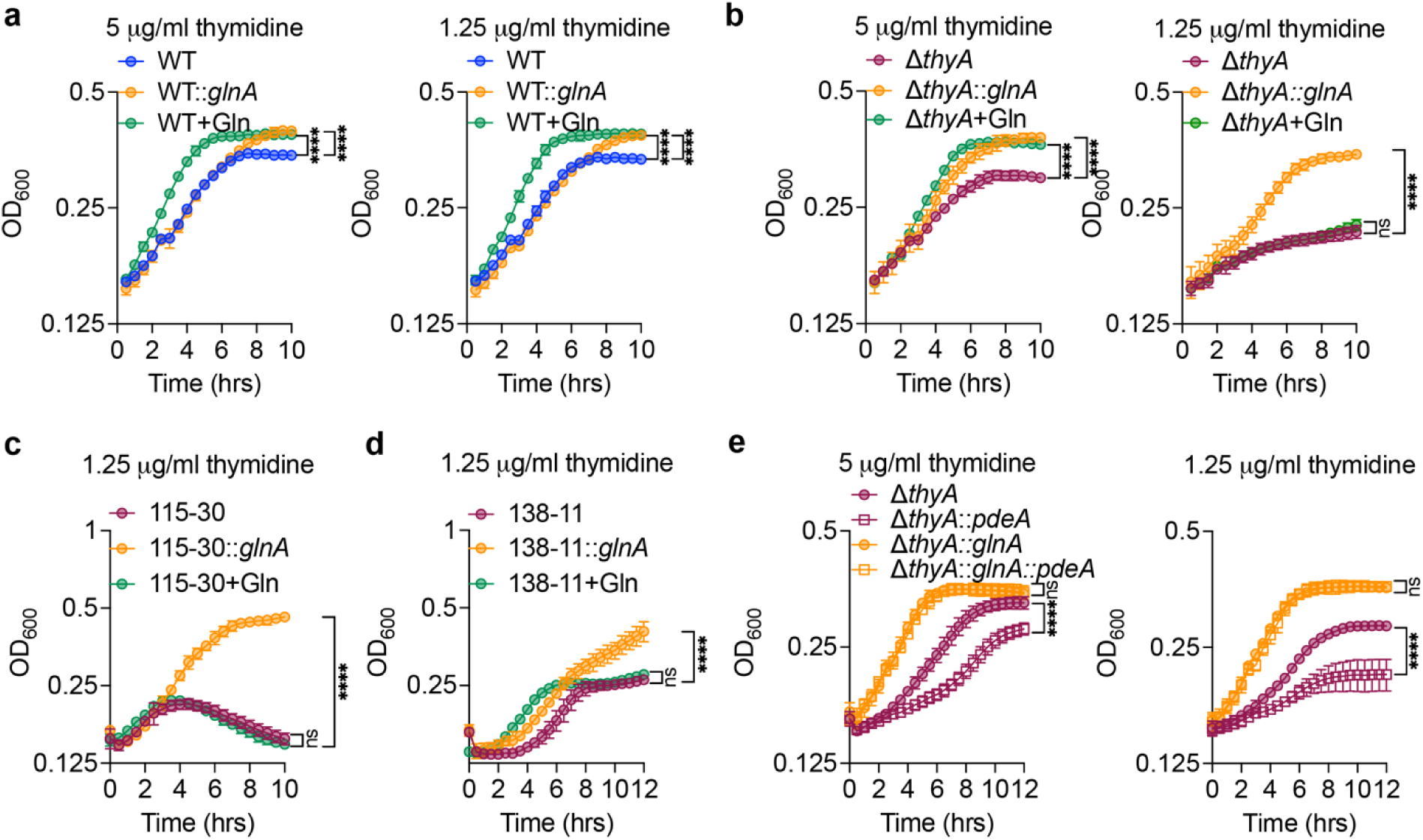
*De novo* glutamine synthesis promotes the growth of *S. aureus* TD-SCV in low thymidine conditions. **a-b**, Growth curve of *S. aureus* strains grown in SCFM with or without 20 mM glutamine supplementation. **c-d**, Growth curve of *S. aureus* TD-SCV clinical isolates, 115-30 and 138-11, grown in SCFM with or without *glnA* overexpression or 20 mM glutamine supplementation. **e**, Growth curves of *S. aureus* Δ*thyA* with overexpression of *pdeA*, *glnA*, or both, grown in SCFM with varying thymidine supplementation. For all panels, mean values of biological triplicates are plotted, and error bars indicate ±SD, *P* values were calculated using Two-way ANOVA analysis based on the final OD_600_ at 12 hours. Asterisks indicate that differences are statistically significant (****, *P* < 0.0001), and “ns” indicates no significant difference.

### De novo glutamine synthesis promotes the survival of *S. aureus* TD-SCV during infection

We have observed that *de novo* glutamine synthesis promotes the growth of Δ*thyA* under low-thymidine conditions *in vitro* (**Fig. 6b**). Our previous study revealed that macrophages do not contain sufficient thymidine to support the growth of *L. monocytogenes* Δ*thyA* strain^39^. We investigated the intercellular survival of *S. aureus* WT, WT::*glnA*, Δ*thyA*, and Δ*thyA*::*glnA* strains in primary bone marrow-derived macrophages (pBMDMs) by gentamicin protection assays. We found that Δ*thyA* exhibited a significant growth defect in macrophages compared to WT, and this defect was fully rescued by *glnA* overexpression, whereas *glnA* overexpression in WT had no impact on WT survival (**Fig. 7a**). Furthermore, C57BL/6J wild-type mice were intranasally infected with these strains at an inoculum of 5×10^8^ CFU per mouse. We found that *glnA* overexpression only moderately promoted the growth of WT in murine lungs; however, the growth defect of Δ*thyA* was rescued by *glnA* overexpression, as determined by CFU recovery at 16 h post-infection (hpi) (**Fig. 7b**). These data support our hypothesis that *de novo* glutamine synthesis is critical for the growth of TD-SCVs during infection and GlnA could be a drug target to control the growth of *S. aureus* TD-SCVs.

**Figure 7.**
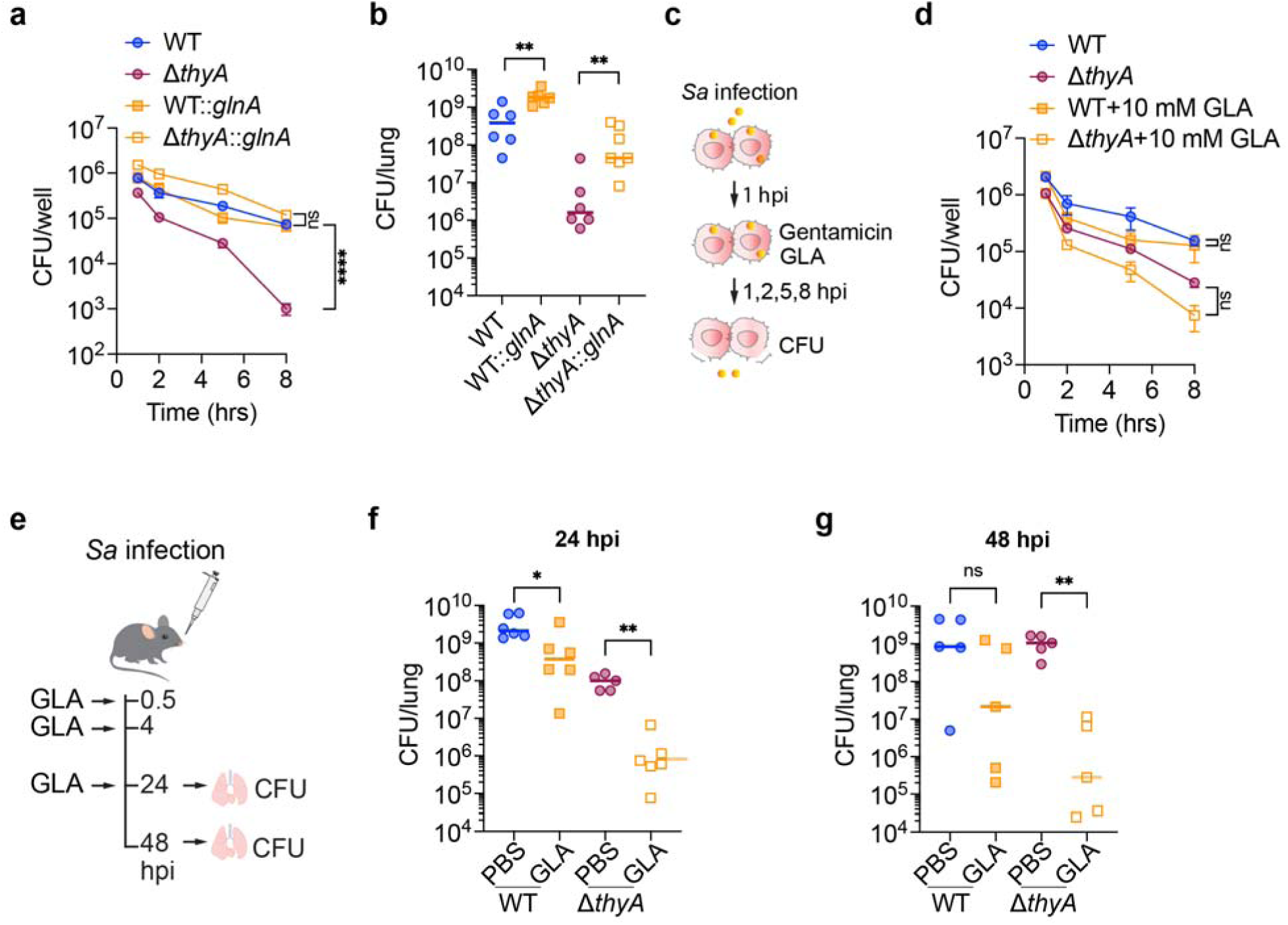
De novo glutamine synthesis promotes the growth of growth of *S. aureus* TD-SCV during infections. **a**, Intracellular growth curves of *S. aureus* WT and Δ*thyA* strains with or without *pdeA* overexpression in pBMDMs. The pBMDMs were infected with *S. aureus* at an MOI of 10. Gentamicin was supplemented at 1 hpi to kill the extracellular bacteria. **b**, CFU recovery of *S. aureus* WT and Δ*thyA* strains with or without *pdeA* overexpression in the lung. The mice were intranasally infected with *S. aureus* at an inoculum of 5 × 10^8^ CFU per mouse. At 16 hpi, the lungs were collected for CFU enumeration. **c**, Schematic of GLA treatment in macrophages infected with *S. aureus*. The pBMDMs were infected with *S. aureus* at an MOI of 10. Gentamicin and GLA were added at 1 hpi. **d**, Intracellular growth curves of *S. aureus* WT and Δ*thyA* strains in pBMDMs as described in **c**. **e**, Schematic of GLA treatment in mice infected with *S. aureus*. The mice were intranasally infected with *S. aureus* at an inoculum of 5 × 10^8^ CFU per mouse. GLA was administered intranasally at 0.5, 4, and 24 hpi. Lungs were harvested at 24 or 48 hpi for CFU enumeration, corresponding to two or three GLA doses, respectively. **f-g**, CFU recovery of *S. aureus* WT and Δ*thyA* strains in the lung at 24 and 48 hpi as described in **e**. For macrophage infection assays, mean values of biological triplicates are plotted, and error bars indicate ±SD, *P* values were calculated using Two-way ANOVA analysis. For all mouse infection panels, biological replicates are plotted. Horizontal black bars indicate the median of the data. *P* values were calculated using Kruskal-Wallis followed up by Uncorrected Dunn’s test. Asterisks indicate that differences are statistically significant (*, *P* < 0.05; **, *P* < 0.01; ***, *P* < 0.001, ****, *P* < 0.0001), and “ns” indicates no significant difference.

To our hypothesis, pBMDMs were infected with WT and Δ*thyA* strains, and the glutamine synthetase inhibitor glufosinate ammonium (GLA) was added at 10 mM at 1 hpi (**Fig. 7c**). We found that GLA slightly inhibited the intracellular growth of the Δ*thyA* strain, whereas it did not inhibit the growth of the WT strain (**Fig. 7d**). To further explore the potential of using GlnA inhibitors for *S. aureus* TD-SCV treatment *in vivo*, C57BL/6J wild-type mice were intranasally infected with WT and Δ*thyA* strains at an inoculum of 5×10^8^ CFU per mouse. GLA was then intranasally instilled at 20 mg/kg at 0.5, 4 hpi, and 24 hpi, and the lungs were harvested at 24 and 48 hpi for CFU enumeration (**Fig. 7e**). We found that although GLA inhibited the growth of both the WT and Δ*thyA* strains in the lungs, it exhibited a significantly greater inhibitory effect against the Δ*thyA* strain, reducing its bacterial burden by approximately 3 logs at 24 hpi (**Fig. 7f**) and 4 logs at 48 hpi (**Fig. 7g**), as determined by CFU enumeration. These data indicate that inhibition of GlnA can significantly suppress the growth of *S. aureus* TD-SCVs in the lungs and has the potential to serve as an adjuvant to antifolate antibiotics for combating antibiotic resistance.

## Discussion

In this study, we identified a previously uncharacterized c-di-AMP-dependent pathway that regulates glutamine synthesis in *S. aureus*, compensating for the absence of the canonical glutamine regulatory proteins TnrA, GlnK, and GlnB. We found that the c-di-AMP receptor PstA interacts with GlnR and cooperates with GlnA to enhance the DNA-binding activity of GlnR, thereby repressing glutamine synthesis genes in response to changes in intracellular c-di-AMP levels. These findings resolve a long-standing question regarding the physiological function of the highly conserved c-di-AMP receptor PstA in Firmicutes. Beyond defining the mechanism of glutamine regulation in *S. aureus*, we demonstrated that *de novo* glutamine synthesis, rather than exogenous glutamine uptake, is essential for the survival of *S. aureus* TD-SCVs during murine lung infection. We further showed that elevated c-di-AMP production sustains glutamine synthesis under thymidine-limited conditions, enabling TD-SCV survival and potentially contributing to the chronic and recurrent infections observed in the CF airway, where thymidine availability is limited. Finally, pharmacological inhibition of glutamine synthesis markedly reduced the pulmonary bacterial burden of *S. aureus* TD-SCVs, identifying glutamine synthesis as a promising therapeutic target for TD-SCV infections. Our findings also suggest that glutamine synthetase inhibitors could serve as effective adjuvants to antifolate therapy to overcome antifolate resistance in *S. aureus* and other bacterial pathogens.

Glutamine is essential for protein synthesis, serves as a nitrogen donor for nucleic acid biosynthesis and diverse transamidation reactions, and also functions as a signaling molecule in bacteria^21^. *S. aureus* is a human pathogen capable of infecting multiple organs in both extracellular and intracellular niches and must rapidly adjust its metabolism to adapt to nitrogen starvation and antibiotic stress^40,41^. However, the regulatory mechanisms governing glutamine metabolism in *S. aureus* remain largely unknown. In this study, we address this knowledge gap by characterizing the interplay between c-di-AMP signaling and glutamine metabolism. A striking feature of *S. aureus* glutamine metabolism is that glutamine acts as the amide donor for the amidation of the peptidoglycan (PG) precursor Lipid II^42^. Reduced intracellular glutamine levels result in decreased PG amidation and diminished PG cross-linking by transpeptidases, which in turn reduces resistance to β-lactam antibiotics and increases sensitivity to lysozyme^43^. Interestingly, clinical isolates with elevated c-di-AMP concentrations, which is attributed to inactivating mutations in the phosphodiesterase GdpP, exhibit next-generation β-lactam resistance in strains lacking PBP2a-mediated β-lactam resistance^44–47^. These data indicate that c-di-AMP-mediated regulation of glutamine metabolism may also play a role in β-lactam resistance in *S. aureus*, underscoring the clinical importance of characterizing this regulatory pathway.

*S. aureus* TD-SCVs exhibit hypermutability and multidrug resistance^38^, which may facilitate their ability to evade the host immune system, contribute to chronic and recurrent infections, and develop drug- or multidrug-resistant strains. Our study reveals a previously overlooked finding: although exogenous glutamine is available in lung sputum and within host cells, *S. aureus* TD-SCVs exhibit diminished glutamine uptake and instead rely on *de novo* glutamine synthesis for survival. Furthermore, bacteria use glutamine as a substrate for pyrimidine synthesis. *S. aureus* TD-SCV grown under low thymidine conditions exhibits elevated pyrimidine synthesis; however, the absence of the ThyA enzyme, which converts dUMP to dTMP, leads to the accumulation of dUTP. dUTP can be incorporated into DNA, helping to sustain DNA replication and facilitate bacterial growth; at the same time, this process promotes mutations through the action of DNA repair pathways. Therefore, our data also provide a mechanistic explanation for why S*. aureus* TD-SCVs are hypermutable.

PII signaling proteins are multitasking regulators found across all domains of life. They modulate cellular processes, including nitrogen metabolism, through protein-protein interactions that are altered upon metabolite binding^32^. PstA is conserved in Firmicutes and is distinct from other PII signaling proteins in that it does not bind nitrogen- or carbon-related metabolites but instead binds the second messenger c-di-AMP^33,34,48^. C-di-AMP is essential for the growth of many Firmicutes under certain growth conditions, including *B. subtilis*^49^, *S. aureus*^50^, *Listeria monocytogenes*^51^, and *E. faecalis*^52^. However, the function of PstA, as well as how it is regulated by c-di-AMP, has remained a longstanding question. We previously showed that PstA promotes thymineless death in *L. monocytogenes* during infection, a process inhibited by elevated intracellular c-di-AMP^39^. Here, we demonstrate that the c-di-AMP-PstA pathway regulates *S. aureus* glutamine synthesis in response to glutamine availability, which in turn regulates the survival of *S. aureus* TD-SCVs during lung infection. Our study not only clarifies the regulatory role of PstA in central metabolism with bacterial pathogenesis, but also underscores the functional versatility of PstA. The multitasking nature of the PII family is attributed to their ability to interact with different target proteins in response to binding distinct ligands or undergoing post-translational modifications (PTMs)^32^; however, such mechanisms remain poorly characterized for PstA. Future studies focusing on characterizing PstA PTMs under different c- di-AMP concentrations may uncover additional regulatory functions.

Our study provides compelling evidence that PstA interacts with GlnR and enhances its DNA-binding activity, thereby identifying GlnR as the first protein target of PstA in bacteria. However, we did not observe a supershift in EMSA assays when characterizing PstA-mediated enhancement of GlnR DNA binding. Interestingly, a similar phenotype has been observed for GlnA-mediated promotion of GlnR DNA-binding activity in our study and others^23^. Together, these data indicate that PstA likely interacts with GlnR transiently rather than forming a stable complex. Such a transient interaction may be advantageous for bacteria, as it would enable more sensitive and efficient regulation in response to fluctuating c-di-AMP levels. In addition, although our data clearly demonstrate that PstA act cooperatively with GlnA to enhance DNA-binding activity of GlnR, we did not detect direct binding between PstA and GlnA alone, nor any impact on GlnA’s enzymatic activity (**Extended Data Fig. 5**). PstA may not directly interact with GlnA, or this interaction may require the presence of both GlnR and DNA. Future structural studies of the GlnA-GlnR-PstA-DNA complex may provide insights into the precise molecular mechanisms involved.

Taken together, by elucidating how c-di-AMP regulates *de novo* glutamine synthesis and the survival of *S. aureus* TD-SCVs, our study will, for the first time, reveal the intricate interplay among glutamine, thymidine, and c-di-AMP in controlling *S. aureus* TD-SCV growth in CF sputum and during infection. In doing so, this work addresses a long-standing question underlying persistent *S. aureus* TD-SCV infections and identifies glutamine synthesis as potential therapeutic targets for drug-resistance *S. aureus* TD-SCV treatment.

## Methods

### Bacterial growth conditions

*S. aureus* Newman and RN4220 strains were grown in brain heart infusion broth (BHI) and incubated at 37 °C. *Escherichia coli* used for cloning was grown in Lysogeny broth (LB) at 37 °C. When necessary, relevant antibiotics were added to the cultures with the following final concentration: 1 μg/mL anhydrotetracycline (Antet), 10 μg/mL chloramphenicol (Cm), and 2 μg/mL erythromycin (Erm). The bacterial strains and plasmids used in this study are listed in **Supplementary Table 1**.

### Bacterial genetic manipulation

The *S. aureus* Newman Δ*thyA* strain was constructed in our previous study^15^*. S. aureus* Newman Δ*pstA* strain were constructed by allelic exchange using pIMAY^53^. Briefly, genomic DNA from *S. aureus* strain Newman was used as a template to generate fragments containing 500 bp upstream and 500 bp downstream of *pstA* (*NWMN_0447*) using the primers listed in **Supplementary Table 2**. The resulting amplimer was cloned into pIMAY and then transformed to *E. coli* DC10B. Allelic exchange was accomplished in two steps. First, the pIMAY_deletion cassette, purified from *E. coli* DC10B, was electroporated into the WT Newman strain and grown on BHI plates containing 10 µg/mL Cm at 30 °C and then at 37 °C to ensure integration. The resulting integration colonies were grown overnight without selection and then plated on BHI plates containing 1 µg/mL Antet to promote allelic exchange. Single colonies were patched onto BHI plates with Antet and BHI plates with Cm, and grown overnight at 37 °C. Cm-sensitive (Cm^S^) and Antet-resistant (Antet^R^) colonies were screened for the desired mutation using colony PCR.

Δ*pstA* complementation strain (Δ*pstA*::*pstA*) were constructed by cloning *pstA* along with its original promoter into pJC1112 using the primers listed in **Supplementary Table 2**. The recombinant pJC1112 vector was electroporated into *S. aureus* strain RN4220 containing pRN7023 (**Supplementary Table 1**), which facilitates single-copy chromosomal integration of constructs into the SaPI1 site by encoding the SaPI1 integrase^54^. Phage φ-11 was propagated in RN4220/pRN7023 to package the pJC1112 vector, and this was followed by phage transduction to introduce the pJC1112 vector into the Δ*pstA* strain. Plasmid integration was selected using 2 μg/mL Erm and confirmed by PCR.

The *glnA* overexpression strains were constructed by cloning *S. aureus glnA* (*NWMN_1217*) into the pEPSA5 plasmid^55^, and *pdeA* overexpression strains were constructed by cloning the *L. monocytogenes pdeA* gene (*LMRG_02481*) into the pBAV1K-E* plasmid^56^. The recombinant pEPSA5 or pBAV1K-E* vectors were passaged in *E. coli* DC10B and then electroporated into *S. aureus* strains. The resulting *glnA* overexpression strains were grown in BHI containing 10 μg/mL Cm, and *glnA* expression was induced by supplementation with 2% xylose. The *pdeA* overexpression strains were grown in BHI containing 100 μg/mL kanamycin, and *pdeA* expression was induced by the addition of 2 mM theophylline.

### Bacterial growth curves assays

Bacterial growth curve assays were performed in a 96-well cell suspension plate using Microplate Reader (BioTek). An overnight culture of *S. aureus* grown in SCFM^57^ or BHI at 37°C with shaking was washed twice with phosphate-buffered saline (PBS) and pelleted by centrifugation. The *S. aureus* pellet was resuspended in SCFM to an OD_600_ of 0.05 with various nitrogen sources. A 200 μL aliquot of the bacterial suspension was added to a 96-well suspension plate, sealed with oxygen-permeable film, and cultured at 37°C with shaking. The OD_600_ was measured every 30 minutes using a Microplate Reader (BioTek).

### 3H-glutamine uptake assays

Both WT and Δ*thyA S. aureus* Newman strains were grown overnight in BHI supplemented with 5 μg/mL thymidine at 37°C with shaking. The bacteria were then washed twice with PBS and resuspended in SCFM to an OD_600_ of 0.05 with various concentrations of thymidine. The bacterial suspensions were supplemented with either 0.5 μCi ^3^H-glutamine alone or 0.5 μCi ^3^H-glutamine plus 20 mM glutamine, and grown at 37 °C with shaking until mid-log phase. The bacterial pellets were collected by centrifugation, washed three times with PBS, resuspended in 20 μL of H_2_O, and transferred to scintillation vials prefilled with 20 mL of liquid scintillation cocktail (BD). The ^3^H signal was then measured using a liquid scintillation counter.

### RNA isolation and qRT-PCR

A total of 5 mL of *S. aureus* grown in SCFM was collected by centrifugation, flash-frozen in liquid nitrogen, and stored at -80°C for 2 hours before RNA isolation. The bacterial pellet was resuspended in 400 µL AE buffer (50 mM NaOAc, pH 5.2; 10 mM EDTA), then mixed with 80 µL of 10% SDS and 500 µL of acidified phenol:chloroform (pH 4.5). The mixture was vortexed at 2,000 rpm for 20 minutes and centrifuged for 30 minutes at 4°C. The aqueous layer containing the RNA was then transferred to a tube containing 50 µL of 3M NaOAc (pH 5.2) and 1.0 mL of 100% ethanol. The samples were incubated at -20°C for 1 hour and then centrifuged to precipitate the RNA. The RNA pellet was washed with 500 µL of 70% ethanol and dissolved in nuclease-free water. A total of 1 µg of RNA was used for cDNA synthesis with Maxima H Minus Reverse Transcriptase with DNase (Fisher Scientific), following the manufacturer’s instructions. The qRT-PCR was performed using the resulting cDNA template, primers listed in **Table S2**, and SYBR™ Green Master Mix (ABI). The relative expression of genes was calculated using the 2^-ΔΔCt^ method, with 16S rRNA as the internal control.

### Promoter activity assay

The promoter region of the *glnRA* operon was amplified from *S. aureus* Newman genomic DNA and fused with the coding sequence of the *gfp* gene using the primers listed in **Supplementary Table 2**. The construct was cloned into pEPSA5 and first electroporated into *S. aureus* RN4220, and subsequently introduced into *S. aureus* Newman strains. For promoter activity measurements, overnight cultures grown in BHI medium supplemented with 10 µg/mL Cm were back-diluted to an OD_600_ of 0.05 into fresh BHI medium with different concentrations of glutamine supplementation and transferred to 96-well suspension culture plates. Plates were sealed with oxygen-permeable film and incubated at 37°C with shaking for 16 h. Following incubation, cultures were washed and resuspended in PBS and transferred to a fresh plate for analysis. Optical density (OD_600_) and GFP fluorescence (excitation: 458 nm; emission: 530 nm) were measured using a BioTek microplate reader. Promoter activity was quantified as fluorescence intensity normalized to OD_600_.

### C-di-AMP quantification

C-di-AMP levels were quantified as previously described by the Gründling Lab using a pCN34e plasmid encoding YFP under the control of the c-di-AMP-binding riboswitch kimA^36^. *S. aureus* strains containing the riboswitch biosensor were grown overnight in BHI medium supplemented with 10 µg/mL erythromycin and then back-diluted to an OD_600_ of 0.05 into fresh BHI medium containing varying concentrations of L-glutamine. Cultures were transferred to 96-well suspension culture plates, sealed with oxygen-permeable film, and incubated at 37 °C with shaking for 16 h. Following incubation, cultures were washed and resuspended in PBS and transferred to a fresh plate for analysis. OD_600_ and YFP fluorescence (excitation: 488 nm; emission: 535 nm) were measured using a BioTek microplate reader. C-di-AMP levels were quantified as fluorescence intensity normalized to OD_600_.

### Bacterial whole cell proteomics

A total of 100 mL of mid-log culture of *S. aureus* Newman WT and Δ*thyA*, grown in the presence of 5 or 1.25 μg/mL of thymidine, was collected by centrifugation and washed twice with PBS. The bacterial pellet was resuspended in 1 mL of lysis buffer (8M urea, 75 mM NaCl, 50 mM Tris, 0.05% Rapigest, pH 8.2) and lysed using sonication. The soluble lysates were collected by centrifugation, and DTT was added to a final concentration of 5 mM, followed by incubation for 30 minutes at 56°C to reduce disulfide bonds. The proteins in the lysate were alkylated by adding 14 mM iodoacetamide and incubating for 30 minutes in the dark at room temperature. Subsequently, the iodoacetamide was quenched by adding an additional 5 mM DTT, followed by a 15-minute incubation in the dark. The protein mixture was then diluted 1:5 in 25 mM Tris (pH 8.2) to reduce the urea concentration to 1.6 M, and CaCl_2_ was added to a final concentration of 1 mM, both of which are necessary for efficient trypsin activity. Trypsin was added at a 1:200 trypsin:protein (w/w) ratio for overnight digestion at 37°C. The sample was then cooled to room temperature, and 0.4% trifluoroacetic acid was added to ensure the pH was <2.0. The digested protein was purified using a C18 column (Nest Group) following the manufacturer’s protocol, and the peptides were resuspended in 0.1% formic acid. The peptides were analyzed using an Easy-nLC 1000 liquid chromatograph coupled with an Orbitrap Eclipse mass spectrometer (Thermo Scientific). Raw spectral data were processed using the MaxQuant software, and peptide sequences were searched against the *S. aureus* Newman reference proteome (UniProt). The relative abundance of proteins was determined using the label-free quantification (LFQ) intensity metric generated by the MaxLFQ algorithm^58,59^.

### Protein expression and purification

All His-tagged proteins were expressed from pET vectors in *E. coli* BL21(DE3) strain and purified using Ni-affinity and size-exclusion chromatography. In brief, *E. coli* BL21(DE3) was transformed with the expression constructs and grown in LB broth at 37°C to an OD_600_ of 0.5-0.8 before induction with 1 mM isopropyl-β-D-galactopyranoside (IPTG) at 16°C overnight. Harvested cell pellets from 1 L cultures were resuspended in 30 mL of ice-cold buffer A (30 mM K_2_HPO_4_, 300 mM NaCl, pH 8.0) containing 1 mM PMSF and 20 mM imidazole, lysed by sonication, and centrifuged at 15,000g for 45 minutes at 4°C. The supernatants were applied to

0.5 mL of Ni-NTA resin, washed with 100 mL of buffer A containing 20 mM imidazole, and eluted with 5 mL of buffer A containing 250 mM imidazole. The elution was subsequently purified using a HiPrep 26/70 Sephacryl S200 column pre-equilibrated with a storage buffer on a Bio-Rad FPLC system. The storage buffer for PstA and GlnA was PBS, while for GlnR, it was 30 mM K_2_HPO_4_, 300 mM NaCl, pH 8.0. Eluted proteins were concentrated, examined by SDS-PAGE, and quantified using a NanoDrop based on the extinction coefficient or by the Bradford assay (Bio-Rad). The purified protein was flash-frozen in 100 μL aliquots using liquid nitrogen and stored at –80°C.

### Electrophoretic mobility shift assay (EMSA)

The 5’-FAM-labeled positive-sense DNA strand and the unlabeled negative-sense strand (**Supplementary Table 1**) were synthesized by Integrated DNA Technologies (IDT) and annealed by incubating them at 95 °C for 5 minutes, followed by cooling to room temperature. The DNA probes were then incubated with different amounts of GlnR proteins in EMSA buffer (50 mM Tris-HCl, pH 8.0; 2.5 mM MgCl_2_; 1 mM DTT; 100 mM KCl; and 10% glycerol) at 25 °C for 20 minutes. To investigate the effects of GlnA and PstA on the DNA binding of GlnR, different amounts of these proteins were added as indicated. Glutamine, glutamic acid, and (NH_4_)_2_SO_4_ were prepared in aqueous solution and adjusted to pH 8.0 before supplementation. The mixtures were subsequently subjected to 6% native polyacrylamide gel electrophoresis at 150 V for 50 minutes on ice. Gel images were obtained using an Azure Sapphire Imager.

### Differential radial capillary action of ligand assay (DRaCALA)

For DNA binding activity of GlnR, the 5’-FAM-labeled DNA probes (**Supplementary Table 2**) were incubated with different amounts of GlnR protein in 50 mM Tris-HCl, pH 8.0; 2.5 mM MgCl_2_; 1 mM DTT; 100 mM KCl at 25 °C for 20 minutes. The mixtures (4 μL per sample) were subsequently spotted onto a nitrocellulose membrane (0.2 μm), air-dried, and imaged using an Azure Sapphire Imager.

For testing of c-di-AMP binding ability of various proteins, 1 µM α-[³²P]ATP was incubated with 10 µM *B. subtilis* diadenylate cyclase DisA in DisA buffer (40 mM Tris-HCl, pH 7.5; 100 mM NaCl; 20 mM MgCl_2_) at 30 °C overnight. The resulting ³²P-c-di-AMP was then captured by incubating the reaction with 10 µM RECON^60^ protein conjugated to Ni-NTA beads for 30 minutes and washed at least three times. The beads were then incubated at 95 °C for 5 minutes to release ³²P-c-di-AMP. The purity of ^32^P-c-di-AMP was assessed using Polygram CEL300 PEI TLC plates (Macherey-Nagel) in a buffer containing a 1:1.5 (vol/vol) ratio of saturated (NH_4_)_2_SO_4_ and 1.5 M NaH_2_PO_4_ (pH 3.6). The tested proteins were mixed with 1 μL of ^32^P-c-di-AMP, and DisA binding buffer was added to bring the final volume to 10 μL. The mixture was incubated for 10 minutes at room temperature. Then, 4 μL of the mixtures were spotted onto a nitrocellulose membrane (0.2 μm), air-dried, exposed to a phosphor screen, and imaged using an Azure Sapphire Phosphor Imager.

### Biolayer interferometry (BLI) assays

The binding kinetics of PstA with GlnR and of GlnR with DNA were analyzed using an Octet RED96 system (ForteBio) equipped with Streptavidin biosensors (Sartorius), with the dissociation constant (*K*_D_) calculated from steady-state analysis using Octet Data Analysis Software 9.0.

To test PstA-GlnR interactions, the C-terminal Avi-tagged PstA protein was biotinylated by incubating it with BirA in 0.05 M bicine buffer (pH 8.3) containing 10 mM ATP, 10 mM MgOAc, and 50 μM D-biotin, followed by overnight incubation at 4 °C. For every 10 nmol of PstA (at 40 μM), 2.5 μg of BirA was used. The biotinylated PstA was then purified using a streptavidin mutein matrix (Roche) following the manufacturer’s instructions. The biotinylated PstA was subsequently diluted to 100 nM in PBS containing 0.02% Tween 20 and 1% BSA in a total volume of 200 μL for conjugation with the Streptavidin Biosensor. Various concentrations of GlnR prepared in 20 mM HEPES (pH 8.0), 100 mM KCl, 2.5 mM MgCl_2_, 1 mM DTT, 0.2% BSA, and 0.05% Tween 20 were used to test the interaction with the PstA-conjugated biosensor.

To test the effects of PstA on the DNA-binding ability of GlnR, a 5’-biotinylated DNA probe (GCTTTATGTTAGAAAACCTGACATATTTT) containing the GlnR binding site was diluted to 1 μM in 20 mM HEPES (pH 8.0), 100 mM KCl, 2.5 mM MgCl_2_, 1 mM DTT, 0.2% BSA, and 0.05% Tween 20 in a total volume of 200 μL for conjugation with the Streptavidin Biosensor. Various concentrations of GlnR in the same buffer were used to test the interaction with the DNA-conjugated biosensor in the presence or absence of PstA.

### Macrophage infections

Primary bone marrow-derived macrophages (pBMDMs) were differentiated from the bone marrow of C57BL/6J wild-type mice for 6 days in DMEM GlutaMAX supplemented with 20% FBS, 10% L929 supernatant, and 100 U/mL penicillin-streptomycin. For the study of intracellular survival of *S. aureus*, 0.5×10^6^ pBMDMs were plated in 24-well tissue culture plates and incubated overnight at 37 °C with 5% CO_2_ in BMM medium (DMEM GlutaMAX supplemented with 10% FBS and 10% L929 supernatant). Mid-exponential-phase *S. aureus* cultures grown in BHI containing 10 µg/mL thymidine were washed twice with PBS and resuspended in BMM medium. Macrophages were infected with 500 µL of the *S. aureus* suspension at a multiplicity of infection (MOI) of 10. After a 1-hour incubation, macrophages were washed once with PBS and incubated with fresh BMM medium containing 100 µg/mL gentamicin to eliminate extracellular bacteria. At the indicated time points post-infection, macrophages were washed once with PBS and lysed by the addition of H_2_O, followed by incubation at 37 °C for 10 minutes. Appropriate dilutions of the lysates were plated on BHI agar supplemented with 10 µg/mL thymidine and incubated overnight at 37 °C for *S. aureus* CFU enumeration.

### Mouse infections

All the mice used in this study were of the C57BL/6J wild type and purchased from Jackson Laboratories. The mice were maintained under SPF conditions, as ensured by the rodent health monitoring program overseen by the Animal Care Facility at the University of Texas at Arlington. All experiments involving mice were performed in compliance with the guidelines set by the American Association for Laboratory Animal Science (AALAS) and were approved by the Institutional Animal Care and Use Committee (IACUC) at UT Arlington. All experiments were carried out using mice aged 7-8 weeks, matched by gender, age, and body weight.

Prior to animal experiments, *S. aureus* strains were cultured overnight with shaking at 37 °C in BHI, then back-diluted into fresh BHI and grown for 1-2 hours at 37 °C with shaking to reach mid-exponential phase. The bacterial pellets were collected by centrifugation and washed twice with cold PBS, then the suspension was re-suspended with PBS. A 30 μL aliquot of the *S. aureus* suspension was instilled intranasally into mice at an inoculum of 5×10^8^ CFU per mouse under anesthesia. For GLA treatment, mice were intranasally instilled with GLA (20 mg/kg in 30 μL PBS) at 0.5, 4, and 24 hpi. Control mice received 30 μL PBS at the same time points. At indicated time points, mice were euthanized by CO_2_. The lungs were collected, homogenized, and lysed in 5 mL of lysis buffer (0.1% IPEGAL in water) using a Tissue Tearor homogenizer. Bacterial burdens were enumerated by plating serial dilutions on BHI plates. Thymidine (10 μg/mL) was supplemented to support the growth of Δ*thyA* mutants in both BHI broth and plates.

## Acknowledgments

We acknowledge Dr. Joshua J. Woodward of the University of Washington (UW) for his intellectual contributions to this work and for generously sharing *S. aureus* strains. We thank Dr. Lucas R. Hoffman of the UW for providing the clinical *S. aureus* TS-SCV isolates used in this study. We thank the Boutte, Pellegrino, and Ghose laboratories at the University of Texas at Arlington (UTA) for sharing reagents and equipment. We also thank Christine Safieddine of the Research Office Infrastructure of UTA for her assistance with the animal studies. This work was supported by the National Institute of General Medical Sciences (grant 1R35GM162142-01) and National Institute of Allergy and Infectious Diseases (grant 1R01AI198349-01).

## Author Contributions

Q.T. and J.P.L. designed and performed the research experiments, analyzed the data, and wrote the manuscript. Q.T., J.P.L., P.P.T., P.G., and D.R.P. performed the research. S.K. and Y.P. conducted the BLI analyses. P.G. and O.V.E. assisted with tissue culture and animal studies. S.K. and C.C.B. designed the experiments and revised the manuscript. D.J.W. contributed clinical strains. All authors reviewed the final manuscript.

## Competing interests

The authors declare no competing interests.

## Notes

### Competing Interest Statement

The authors have declared no competing interest.

