## Supplemental Figures for "A c-di-AMP-controlled glutamine synthesis pathway promotes persistence of *Staphylococcus aureus* thymidine-dependent small colony variants in the lung"

**Extended Data**

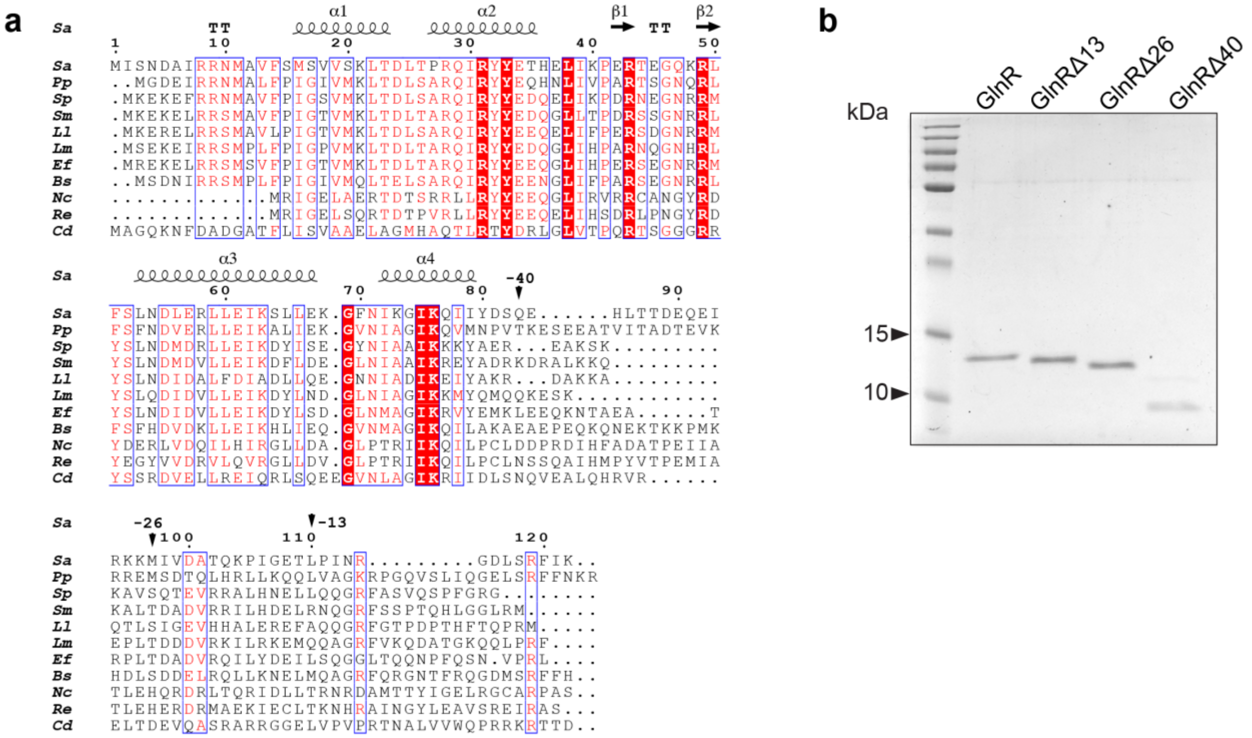


**Extended Data Fig. 1 Expression of GlnR and its C-terminal truncated mutants.** (A) Multiple sequence alignment of GlnR homologs. White letters shaded in red denote identical residues, red letters with white background are similar residues, black letters indicate variable residues and dots represent gaps. The amino acid sequences are from *Staphylococcus aureus* (Accession No.: BAF67488.1), *Paenibacillus polymyxa*, (Accession No.: BED56015.1), *Streptococcus pneumoniae* (Accession No.: EMY89138.1), *Streptococcus mutans* (Accession No.: VTY49302.1), *Lactococcus lactis* (Accession No.: SPS10687.1), *Listeria monocytogenes* (Accession No.: CAC99376.1), *Enterococcus faecalis* (Accession No.: CWW53777.1), *Bacillus subtilis* (Accession No.: WOB00840.1), *Nocardia cerradoensis* (Accession No.:OXR45568.1), *Rhodococcus erythropolis* (Accession No.: OFV75521.1), *Clostridioides difficile* (Accession No.: VTR08026.1). The secondary structure depiction from PDB: 7TEA is shown at the top. The black arrows indicate the site of truncation. (B) SDS-PAGE of GlnR and its C-terminal truncated mutants purified by Ni-NTA resin and size-exclusion chromatography.

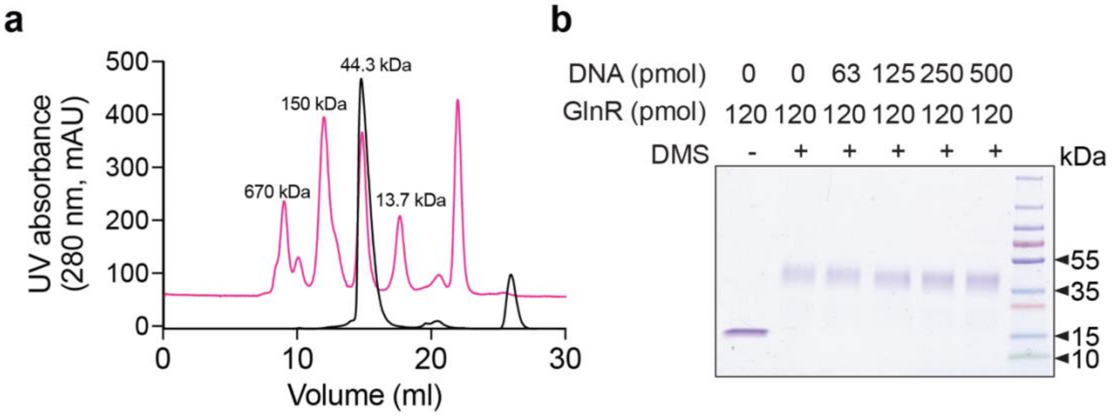


**Extended Data Fig. 2** ***In vitro*** **analysis of GlnR polymerization.** **a**, Size-exclusion chromatography of GlnR protein. The magenta trace represents the protein standard (Sigma), and the black trace represents the GlnR protein analyzed using a Superdex 75 Increase column, respectively. **b**, SDS-PAGE analysis of GlnR polymerization in the presence or absence of *glnAR* probes. GlnR and *glnAR* probes were incubated for 15 min at room temperature before crosslinking with dimethyl suberimidate (DMS) and subsequent SDS-PAGE analysis.

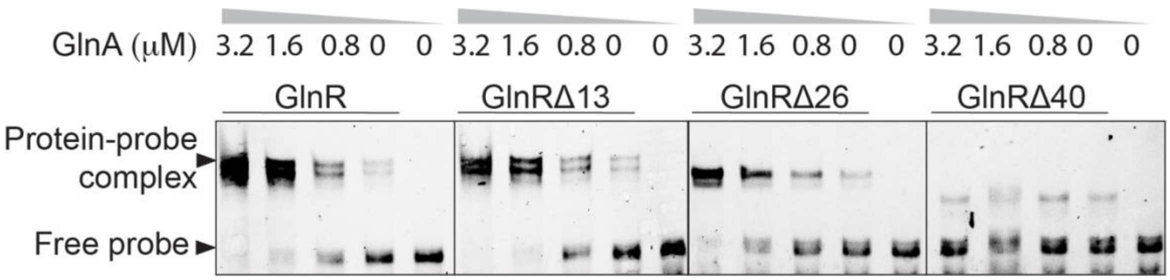


**Extended Data Fig. 3.** EMSA analysis of GlnR and its truncated mutants binding to DNA in the presence of increasing concentrations of GlnA. 0.2 μM 5'-FAM-DNA probe (*glnRA* promoter region) was used.

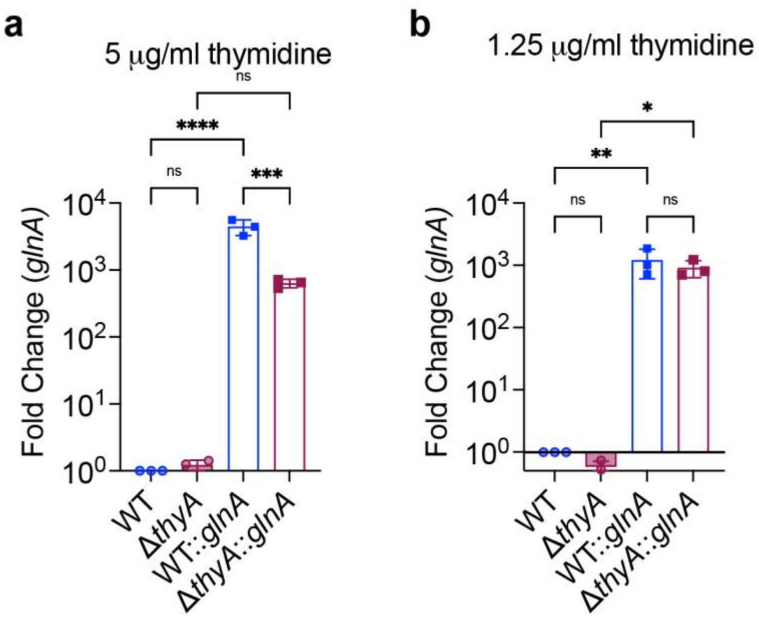


**Extended Data Fig. 4. The expression of *glnA* gene.** Expression of *glnA in* WT, WT::*glnA*, Δ*thyA* and Δ*thyA*::*glnA* strains grown in SCFM in 5 (A) or 1.25 (B) μg/mL thymidine as quantified by qRT-PCR.

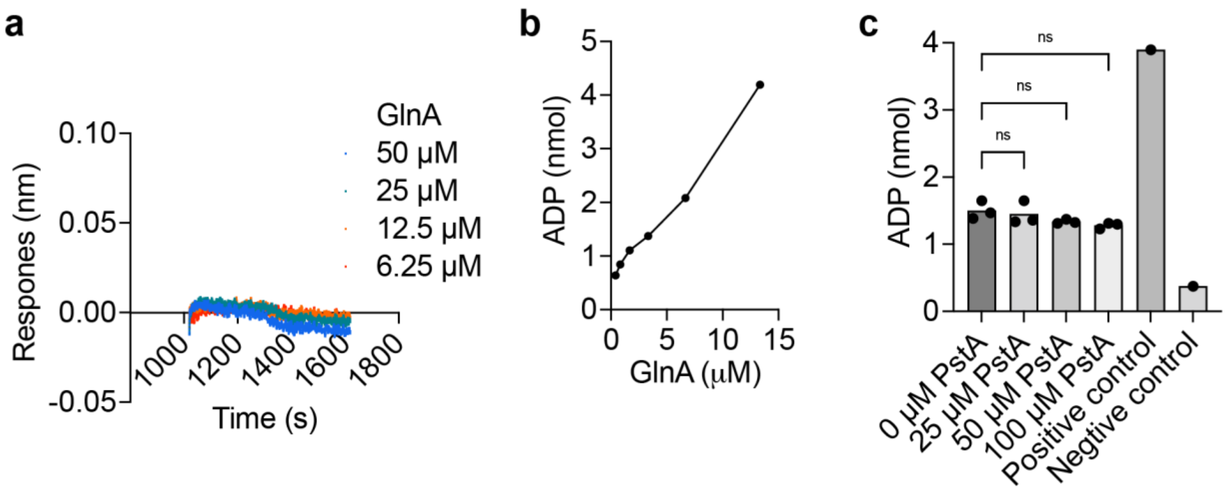


**Extended Data Fig. 5. GlnA enzymatic activity**. **a**, *In vitro* binding of GlnA to PstA analyzed by BLI. Biotinylated-PstA was pre-conjugated with the streptavidin sensor, and an increasing concentration of the GlnA was tested for protein-protein interactions. **b**, Glutamine synthetase (GS) activity at different concentrations of GlnA. **c**, GS activity of GlnA (5 μM) in the presence of increasing concentrations of PstA. GS activity was measured using a Glutamine Synthetase Activity Assay Kit (Abcam, cat#ab284572 ), which quantifies ADP production by glutamine synthetase using glutamate and ATP as substrates. The positive control measured ADP production by a commercial glutamine synthetase preparation from the kit, whereas the negative control measured background signal in the absence of added protein.
