## Supplemental Tables for "A c-di-AMP-controlled glutamine synthesis pathway promotes persistence of *Staphylococcus aureus* thymidine-dependent small colony variants in the lung"

### Supplemental Table 1. Strains used in this study

| Strain# | Strain name | Background | Plasmid | Reference |
|---|---|---|---|---|
| **Plasmids** | | | | |
| TQ752 | DH5a/pCN34e-*yfp* | *E. coli* DH5a | pCN34e-*yfp* | [1] |
| TQ753 | DH5a/pCN34e-kimA-*yfp* | *E. coli* DH5a | pCN34e-kimA-*yfp* | [1] |
| TQ2 | XL1b/pIMAY | *E. coli* XL1Blue | pIMAY | [2] |
| TQ309 | XL1b/PJC1112 | *E. coli* XL1Blue | pJC1112 | [3] |
| TQ139 | DH5a/pEPSA5 | *E. coli* DH5a | pEPSA5 | [4] |
| TQ220 | XL1b/pBAV1K-E* | *E. coli* XL1Blue | pBAV1KE* | [5] |
| TQ307 | RN4220/pRN7023 | *S. aureus* RN4220 | pRN7023 | [6] |
| TQ183 | XL1b/pET20b | *E. coli* XL1Blue | pET20b | Novogen |
| TQ214 | XL1b/pET24b | *E. coli* XL1Blue | pET24b | Novogen |
| TQ213 | XL1b/pET28b | *E. coli* XL1Blue | pET28b | Novogen |
| **Protein expression strains** | | | | |
| TQ377 | $His_6$-GlnR | *E. coli* BL21(DE3) | pET24b-*glnR* | This study |
| TQ391 | $His_6$-GlnRΔ13 | *E. coli* BL21(DE3) | pET24b-*glnR*Δ13 | This study |
| TQ395 | $His_6$-GlnRΔ26 | *E. coli* LOBSTR | pET24b-*glnR*Δ26 | This study |
| TQ392 | $His_6$-GlnRΔ40 | *E. coli* BL21(DE3) | pET24b-*glnR*Δ40 | This study |
| TQ343 | GlnA-$His_6$ | *E. coli* BL21(DE3) | pET24b-*glnA* | This study |
| TQ11 | $His_6$-PstA | *E. coli* BL21(DE3) | pET28b-*pstA* | This study |
| TQ104 | PstA-$His_6$ | *E. coli* BL21(DE3) | pET20b-*pstA* | This study |
| TQ24 | $His_6$-$PstA_{GGFL\text{-}AAAA}$ | *E. coli* BL21(DE3) | pET28b-$pstA_{GGFL\text{-}AAAA}$ | This study |
| TQ311 | PstA-Avi-$His_6$ | *E. coli* XL1b | pET24b-*pstA*-Avi-$His_6$ | This study |
| ***S. aureus* Newman strains** | | | | |
| TQ35 | WT | WT | N/A | |
| TQ37 | Δ*pstA* | Δ*pstA* | N/A | This study |
| TQ355 | Δ*pstA*::*pstA* | Δ*pstA* | pJC1112-*pstA* | This study |
| TQ738 | WT/pEPSA5-P*glnRA-gfp* | WT | pEPSA5-P*glnRA-gfp* | This study |
| TQ739 | Δ*pstA*/pEPSA5-P*glnRA-gfp* | Δ*pstA* | pEPSA5-P*glnRA-gfp* | This study |
| TQ740 | Δ*pstA*::*pstA*/pEPSA5-P*glnRA-gfp* | Δ*pstA*::*pstA* | pEPSA5-P*glnRA-gfp* | This study |
| TQ39 | Δ*thyA* | Δ*thyA* | N/A | [7] |
| TQ164 | Δ*thyA*/pEPSA5 | Δ*thyA* | pEPSA5 | This study |
| TQ691 | Δ*thyA*/pEPSA5-*glnA* | Δ*thyA* | pEPSA5-*glnA* | This study |
| TQ163 | WT/pEPSA5 | WT | pEPSA5 | This study |
| TQ690 | WT/pEPSA5-*glnA* | WT | pEPSA5-*glnA* | This study |
| TQ722 | Δ*thyA*/pEPSA5+pBAV1K-E* | Δ*thyA* | pEPSA5 pBAV1K-E* | This study |
| TQ723 | Δ*thyA*/pEPSA5-*glnA*+pBAV1K-E* | Δ*thyA* | pEPSA5-*glnA* pBAV1K-E* | This study |
| TQ724 | Δ*thyA*/ pEPSA5+pBAV1K-E*-*pdeA* | Δ*thyA* | pEPSA5 pBAV1K-E*-*pdeA* | This study |

| | | | | |
|---|---|---|---|---|
| TQ725 | Δ*thyA*/pEPSA5-*glnA*+pBAV1K-E*-*pdeA* | Δ*thyA* | pEPSA5-*glnA* pBAV1K-E*-*pdeA* | This study |
| TQ746 | WT/pCN34e-*yfp* | WT | pCN34e-*yfp* | this study |
| TQ769 | WT/pCN34E-kimA-*yfp* | WT | pCN34e-kimA-*yfp* | this study |
| **Clinical isolates and their derivatives** | | | | |
| TQ132 | 115-30 | AMT0115-30 SCV 10C3 | NA | [8] |
| TQ137 | 138-11 | AMT0138-11 SCV 10B3 | NA | [8] |
| TQ764 | 115-30/pEPSA5 | AMT0115-30 SCV 10C3 | pEPSA5 | This study |
| TQ759 | 115-30/pEPSA5-*glnA* | AMT0115-30 SCV 10C3 | pEPSA5-*glnA* | This study |
| TQ757 | 138-11/pEPSA5 | AMT0138-11 SCV 10B3 | pEPSA5 | This study |
| TQ758 | 138-11/pEPSA5-*glnA* | AMT0138-11 SCV 10B3 | pEPSA5-*glnA* | This study |

**Supplemental Table 2. Oligonucleotides used in this study**

| Oligo ID | Oligo Name | Sequence (5' to 3') | Reference |
|---|---|---|---|
| | | **GlnR and its mutant protein expression** | |
| TQ145 | *glnR*-F | CATGCCATGGCGATGATATCGAATGATGCAATCAG | This work |
| TQ146 | *glnR*-R | CCGCTCGAGTTTAATAAATCGGGATAAATCACCAC | This work |
| TQ290 | *glnR*Δ13-R | CCGCTCGAGAGTTTCTCCAATAGGCTTTTG | This work |
| TQ310 | *glnR*Δ26-R | CCGCTCGAGTTACTTTTTTCTTATCTCTTGTTCATCT | This work |
| TQ291 | *glnR*Δ40-R | CCGCTCGAGTGTGAGTCATAAATGATTTGTTTAATC | This work |
| | | **GlnA protein expression** | |
| TQ223 | *glnA*-F | CCCATATGGAGGATTTTAAAATGCCAAAAC | This work |
| TQ224 | *glnA*-R | CCCTCGAGATATTGCTTCATGTACTGATCTC | This work |
| | | **PstA and its mutant protein expression** | |
| TQ62 | *pstA*-F | ATACATATGAAAATGATTATAGCGATCGTACAA | This work |
| TQ63 | *pstA*-R | TTCCTCGAGAAATTGATGGAATGCATCAACT | This work |
| NA | GGFL-a | AAATTGGCAACAACAGCCGCCGCCGCCAGAGCGGG TAATACA | This work |
| NA | GGFL-b | TGTATTACCCGCTCTGGCGGCGGCGGCTGTTGTTGC CAATTT | This work |
| | | ***pstA* knockout** | |
| NA | KO*pstA*-1 | CGAGCTCAAATTGTATTAGAAGGCAATGATATG | This work |
| NA | KO*pstA*-2 | TTGTTAGAAGAGGTGTTATAAAATGTAATTCTATAA TACAATCATCAATT | This work |
| NA | KO*pstA*-3 | TTGTTAGAAGAGGTGTTATAAAATGTAATTCTATAA TACAATCATCAATT | This work |
| NA | KO*pstA*-4 | GGGGTACCATGTACACTTTATTTGTGCTTTC | This work |
| | | **Δ*pstA* complementation** | |
| TQ259 | c*pstA*-F | CGGGATCCATTGGCGTTGAAGAAGTAAGA | This work |
| TQ260 | c*pstA*-R | CGGAATTCTTAAAATTGATGGAATGCATCAACT | This work |
| | | **GlnA overexpression** | |
| TQ465 | OE*glnA*-F | CGAGCTCaacgactggaaggagttttaattaTTGGAGGATTTTAA AATGCCA | This work |
| TQ466 | OE*glnA*-R | GGGGTACCTTAATATTGCTTCATGTACTGATC | This work |
| | | **PdeA overexpression** | |
| TQ139 | *LmpdeA*-F | CGGGGTACCTAACAACAAGATGTCAGGCTATTTTC AAAAACGAA | This work |
| TQ140 | *LmpdeA*-R | CTAGACTAGTTTATGTTTCTCCCTTCCAATACG | This work |
| | | **P*glnRA-gfp* reporter construction** | |
| TQ497 | P*glnRA-gfp*-1 | GTTAATTAACGTTAAAATTGCTGTGACAAGAGCTGT TAA | This work |
| TQ498 | P*glnRA-gfp*-2 | TGAAAAGTTCTTCTCCTTTACTCATTGTTCCTCTCCT CTACTTTTGAACT | This work |
| TQ499 | P*glnRA-gfp*-3 | AGTTCAAAAGTAGAGGAGAGGAACAATGAGTAAA GGAGAAGAACTTTTCA | This work |
| TQ500 | P*glnRA-gfp*-4 | GCTCTAGATTATTTGTATAGTTCATCCATGCCATGT GTAATC | This work |
| | | **EMSA probes** | |
| TQ243 | *glnAR*_FAM-F | 6FAM/ATATCAATGTTTTAAGCTTTATGTTAGAAAAC CTGACATATTTTTGAAATCCTAAAAAA | This work |

| | | | |
|---|---|---|---|
| TQ244 | *glnAR*-R | TTTTTTAGGATTTCAAAAATATGTCAGGTTTTCTAA CATAAAGCTTAAAACATTGATAT | This work |
| NA | *glnAR*_biotin-F | 52-Bio/ ATATCAATGTTTTAAGCTTTATGTTAGAAAACCTGA CATATTTTTGAAATCCTAAAAAA | This work |
| TQ 265 | *amtB*_FAM-F | FAM/GCACATTTTATTCCAAAAGATGTAATAAAACT TAACGCATTTTTGCTTTTTATAAATTGT | This work |
| TQ266 | *amtB*-R | ACAATTTATAAAAAGCAAAAATGCGTTAAGTTTTA TTACATCTTTTGGAATAAAATGTGC | This work |

* Restriction enzyme recognition sites are highlighted in red;

6FAM**,** 6-FAM (Fluorescein) is attached to 5'-end of oligo**;**

52-Bio**,** two biotin groups are sequentially placed on the 5'-end of oligo.
